# Cysteine protease cathepsin B promotes high population density-induced mutagenesis, driving genome evolution and competitive growth in response to the crowding stress

**DOI:** 10.64898/2025.12.22.696062

**Authors:** Bin Yu, Yuji Suehiro, Bryan J. Johnson, Eui-Seung Lee, Dongdong Li, Yawen Huang, Joshua Johnson, Guangshuo Ou, James DeGregori, Shohei Mitani, Ding Xue

## Abstract

In a wide variety of species from invertebrates to mammals, overpopulation has been shown to induce low fertility and high mortality. Although density-dependent population regulation is widespread in the animal kingdom, the underlying molecular mechanisms remain poorly understood. We show here that *C. elegans* animals respond to the crowding stress by secreting CPR-4, a homologue of human cathepsin B cysteine protease, to promote chromosomal DNA damage in germ cells, leading to high density-induced deficiencies that include increased embryonic lethality and larval arrest and decreased brood size. CPR-4 mediates these crowding responses through the insulin-like growth factor receptor DAF-2, multiple components in the insulin signaling pathway, and the SKN-1/Nrf transcription factor. Whole genome sequencing analyses of animals from 10 generations of continual growth in the crowded condition reveal that CPR-4-induced DNA damage produces an average of 2.9 more de novo genome mutations per animal per generation and a 75% increase in mutation rate compared with animals grown in the uncrowded condition. CPR-4-induced mutagenesis also facilitates evolution of the genomes through multi-generational crowding selection, leading to biased mutation distributions towards the intergenic regions over the gene bodies and crowd-inducible growth advantage. Our findings suggest that CPR-4 acts as a crucial crowd-responding factor to induce chromosomal DNA damage, leading to density-dependent deficiencies, increased genome mutation rates, and genome evolution and competitive growth of animals in response to the crowding stress.

---

There has been a long-standing debate regarding how species’ populations are regulated for many years^1–3^. Numerous studies have demonstrated that both extrinsic density-independent mechanisms (e.g., environmental factors) and intrinsic density-dependent regulatory mechanisms are involved in controlling population sizes^3–8^. The hypothesis of density-dependent regulation specifies that high population density would cause increased mortality and reduced fertility^9–13^, which serves as a feedback mechanism to curb population growth when the population size approaches the carrying capacity of the environment. This is thought to be central to maintaining the population dynamics at an equilibrium level^14–17^. This important model has been validated in a wide variety of species from invertebrates to mammals and in both natural populations^9–13,18–21^ and laboratory conditions^22–26^. For example, increased larval mortality and reduced reproduction have been observed in fish grown in high density^22,23,25^. Chicken breeding in crowded conditions also results in decreased egg production and increased mortality compared with that in uncrowded conditions^24^. In mice, increased population density leads to a significant decline in fertility rate and an increase in mortality rate from gestation to weaning in both laboratory and wild conditions^25,26^. Moreover, numerous time series demographic studies spanning 145 countries^27^ and 44 developing nations^28^, and a comprehensive study across 174 countries over 69 years^29^ have reached the consistent conclusion that increased population density appears to be directly associated with decreased fertility rate in humans. Although density-dependent population regulation is widespread in the animal kingdom, the molecular players and mechanisms underlying density-dependent regulation remain unknown. It is known, however, that when high population density becomes a long-term issue, density-dependent regulation serves as a selection pressure to facilitate rapid adaptive changes of animals to compete and survive in the stressed and crowded environment^6,30^. Likewise, the molecular mechanisms driving the adaptive changes are not understood.

In a study of how *C. elegans* animals respond to radiation, a frequently encountered environmental stress, we found unexpectedly that high-density growth of *C. elegans* animals in the absence of radiation resulted in elevated secretion of the CPR-4 protein, a crucial radiation-responding factor^31,32^, in a density-dependent manner (Fig. S1a). CPR-4, a homologue of the human cathepsin B cysteine protease, is secreted in response to ultraviolet (UV) or ionizing irradiation to exert both short-range and long-range effects on unexposed cells or tissues in irradiated animals and on unexposed animals^31,32^, indicating that secreted CPR-4 can mediate both intercellular and inter-animal stress signaling. The effects of secreted CPR-4 on unexposed cells or animals due to radiation, also called radiation-induced bystander effects (RIBEs), are numerous, including chromosomal DNA damage, embryonic lethality, larval arrest, stress response, and altered cell death and cell proliferation^31,32^, supporting a role of CPR-4 as a stress responding factor with broad physiological impacts. We therefore investigated whether CPR-4 plays a role in regulating animals’ response to the crowding stress and found that CPR-4 acts as a crucial density-responding factor to increase chromosomal DNA damage in germ cells, leading to multiple density-dependent deficiencies, increased genome mutation rates, and genome evolution and competitive growth of animals to the crowding stress.

## Results

### Secretion of CPR-4 in response to increased *C. elegans* population density

*elegans* animals secret the CPR-4 cysteine protease in response to radiation^31^, which is transported to other parts of the animal through pseudocoelom, a fluid-filled body cavity, and also released into the culture medium. We found that animals grown in the crowded condition with sufficient foods showed increased CPR-4 secretion in the absence of radiation exposure (Fig. S1a). Specifically, larval stage 4 (L4) P*cpr-4::cpr-4::flag*; *cpr-4(tm3718)* animals, which contain a strong loss-of-function mutation (*tm3718)* in *cpr-4* and a single-copy integrated transgene (P*cpr-4::cpr-4::flag*) that fully rescues the *cpr-4* mutant^31^, were grown in plates with plentiful bacterial supplies at different population densities (ranging from 750 animals/plate to 6000 animals/plate; Fig. S1a). No CPR-4::FLAG secretion was observed in plates with a moderate number of L4 or adult animals (less than 750/plate or 0.25/mm^2^; Fig. S1a, lane 1). We define this animal density as uncrowded or not crowded (NC). In contrast, when the density of L4 or adult animals reached more than 3000/plate (1 animal/mm^2^; Fig. S1a, lanes 3-5), increased secretion of CPR-4::FLAG was observed in a density-dependent manner (Fig. S1a). We define these density conditions as crowded (C). Interestingly, secretion of CPR-4::FLAG in crowded conditions was age-dependent. At the density of approximately 3000/plate, only older larvae (L3 and L4) and adult animals, but not younger larvae (L1 and L2), showed CPR-4 secretion, with the adult animals showing much greater CPR-4 secretion than larval animals (Fig. S1b).

To further analyze CPR-4 secretion by animals in crowded conditions, we collected the crowded conditioned medium (C-CM, see Methods) from plates that contained approximately 1500, 2400, and 5400 L4 larvae and from plates with the same numbers of animals 16 hours after L4, respectively (Fig. 1a; see Methods). In parallel, the same numbers of animals were split equally into four different plates and thus were grown in uncrowded conditions (Fig. 1a). The noncrowded conditioned medium (NC-CM) was collected from these four plates sequentially (see Methods). NC-CM and C-CM with equal amounts of total proteins were analyzed by immunoblotting (Fig. 1b). Secreted CPR-4::FLAG was detected in C-CM, but not in NC-CM (Fig. 1b; Fig. S1c). These results indicate that CPR-4 secretion was specifically elevated under the crowded condition in late larvae and adults.

**Fig. 1.**
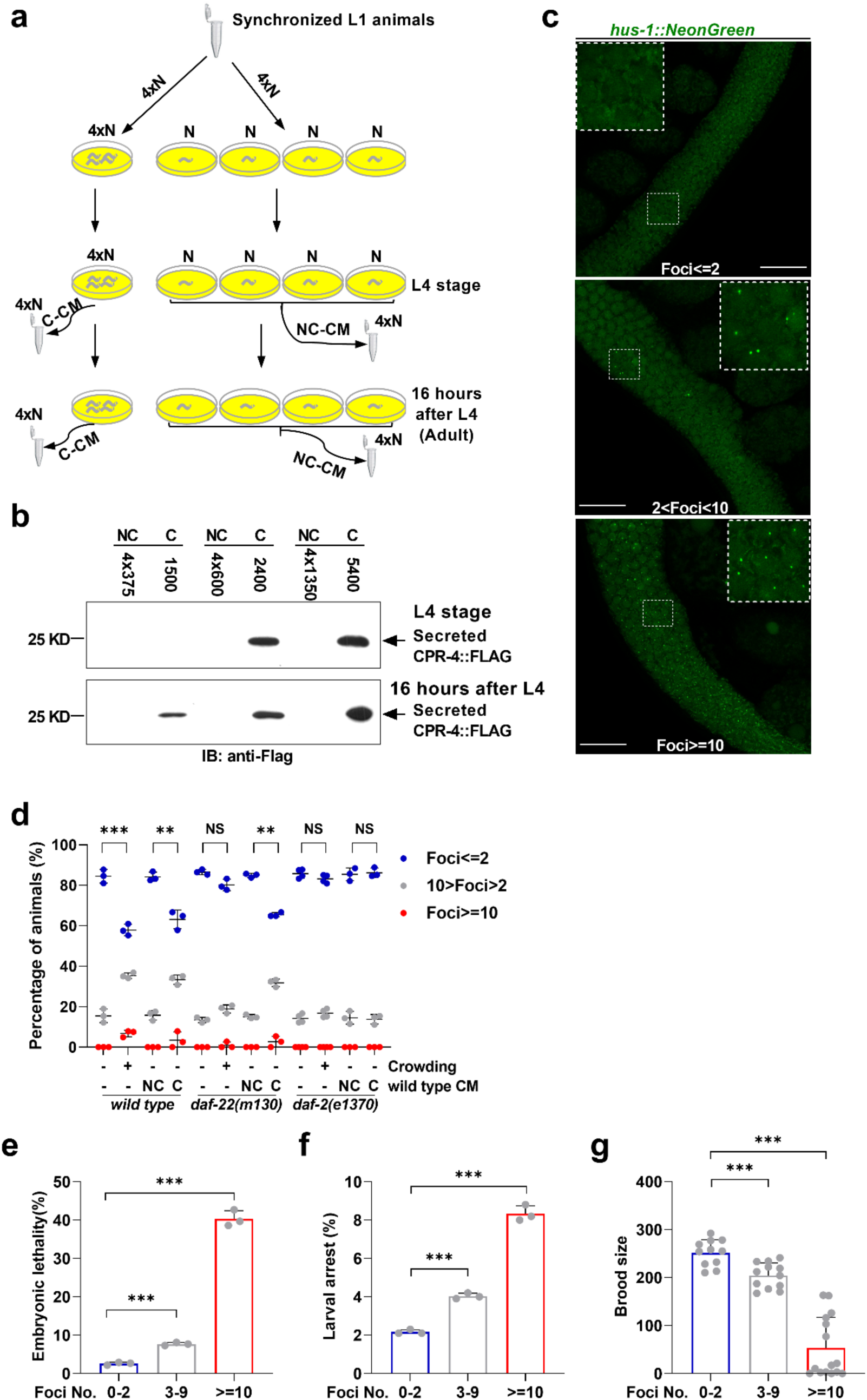
Crowding stress leads to increased secretion of CPR-4, DNA damage in germ cells, and developmental defects in *C. elegans*. **a**, A schematic diagram of the crowding experiments. 4 x N number of synchronized larval stage 1 (L1) animals were grown in one NGM plate (crowded-C) or evenly distributed in four NGM plates (noncrowded-NC). Conditioned medium (CM) was collected when animals reached larval stage 4 (L4) and 16 hours after L4. CM from four noncrowded plates were pooled as NC-CM. **b**, Secreted CPR-4::FLAG proteins were detected in C-CM, but not in NC-CM, derived from P*cpr-4::cpr-4::flag*; *cpr-4(tm3718)* animals grown at high densities. C-CM and NC-CM (6 μg of total proteins each) were resolved by 12% SDS polyacrylamide gel (PAGE) and detected by immunoblotting (IB) using an anti-FLAG monoclonal antibody. **c**, Representative gonad images from *hus-1::neongreen* knockin animals grown in the crowded condition, showing the number of NeonGreen foci ≤2, between 3-9, and ≥10, respectively. Upper left dashed boxes are enlarged images of those highlighted in the middle. Scale bars, 20 μm. **d**, Percentages of *hus-1::neongreen* animals with the indicated genotype and the indicated number of NeonGreen foci in their gonads, grown in uncrowded or crowded conditions, or in uncrowded conditions and then treated with NC-CM or C-CM. **e**-**g**, The percentages of embryonic lethality (**e**) and larval arrest (**f**) observed in progeny and the brood sizes (**g**) of *hus-1::neongreen* adult animals grown in crowded conditions with the indicated number of NeonGreen foci, which were rescued from agarose pads after microscopy screens. Three biologically independent experiments in **d**, **e**, and **f**. The total numbers of adult animals analyzed from left to right are 117, 119, 114, 114, 104, 106, 106, 110, 144, 174, 82, and 80 in **d**, 60, 60, and 54 in **e** and **f**, 11, 12, and 15 in **g**, with the foci numbers from one gonad arm. Total numbers of embryos scored in **e**: 2478 (foci≤2), 1641(foci between 3-9), and 995 (foci≥10). Total numbers of larvae scored in **f**: 2411 (foci≤2), 1514 (foci between 3-9) and 502 (foci≥10). Data are mean ± s.d.. \*\**P* < 0.01, \*\*\**P* < 0.001, two-sided *z*-test in **d** and two-sided, unpaired *t*-test in **e**, **f**, and **g**.

### Elevated CPR-4 secretion causes DNA damage in germ cells

One of the key radiation-induced bystander effects mediated by secreted CPR-4 is genome instability^31,32^, which can be easily detected in unexposed germ cells through a DNA damage marker, the HUS-1 protein^32^. HUS-1 is a DNA damage checkpoint protein and a component of the conserved Rad9, Hus1, and Rad1 complex that localizes to and concentrates at the chromosome damage sites^33^. Using a *hus-1::neongreen* knockin strain^32^, we examined whether increased secretion of CPR-4 due to the crowding stress similarly caused genome instability in germ cells. We found that significantly more HUS-1::NeonGreen foci, indicative of chromosomal DNA damage, were observed in gonads of animals grown in the crowded condition than those grown in the noncrowded condition (Fig. 1c, d). For example, 42% of animals in the crowded condition had more than 2 HUS-1::NeonGreen foci in their gonads and 7% had more than 10 HUS-1::NeonGreen foci, whereas only 15% of animals grown in the uncrowded condition had more than 2 HUS-1::NeonGreen foci and none had more than 10 HUS-1::NeonGreen foci (Fig. 1c, d). Moreover, the extent of DNA damage in germ cells positively correlated with the severity of crowding and the levels of CPR-4 secretion (Fig. S1a, d), with the most germ cell DNA damage observed in animals grown in the highest density. Interestingly, when animals in the uncrowded condition were treated with C-CM or NC-CM for 16 hours, only animals treated with C-CM, but not those treated with NC-CM, had increased numbers of HUS-1::Neongreen foci in their gonads, close to what were observed in animals grown in the crowded condition (Fig. 1d). This indicates that C-CM, but not NC-CM, contained a factor that induced DNA damage and was responsible for the observed increase of germ cell DNA damage. Given that secreted CPR-4::FLAG was present in C-CM, but absent in NC-CM (Fig. 1b) and secreted or recombinant CPR-4 has been shown to be a clastogenic factor that can induce chromosome breakages in normal, unirradiated germ cells non-cell autonomously^32^, we examined if loss of *cpr-4* affected crowding-induced gonad DNA damage. We found that both the *cpr-4(tm3718)* mutation and a deletion removing the entire *cpr-4* coding region, *cpr-4(sm931)*, blocked increased DNA damage induced by crowding, which was restored by the single copy P*cpr-4::cpr-4::flag* transgene (Fig. 2a; Fig S2a). Taken together, these results indicate that secretion of CPR-4 induced by the crowding stress is responsible for the increased DNA damage in *C. elegans* germ cells.

**Fig. 2.**
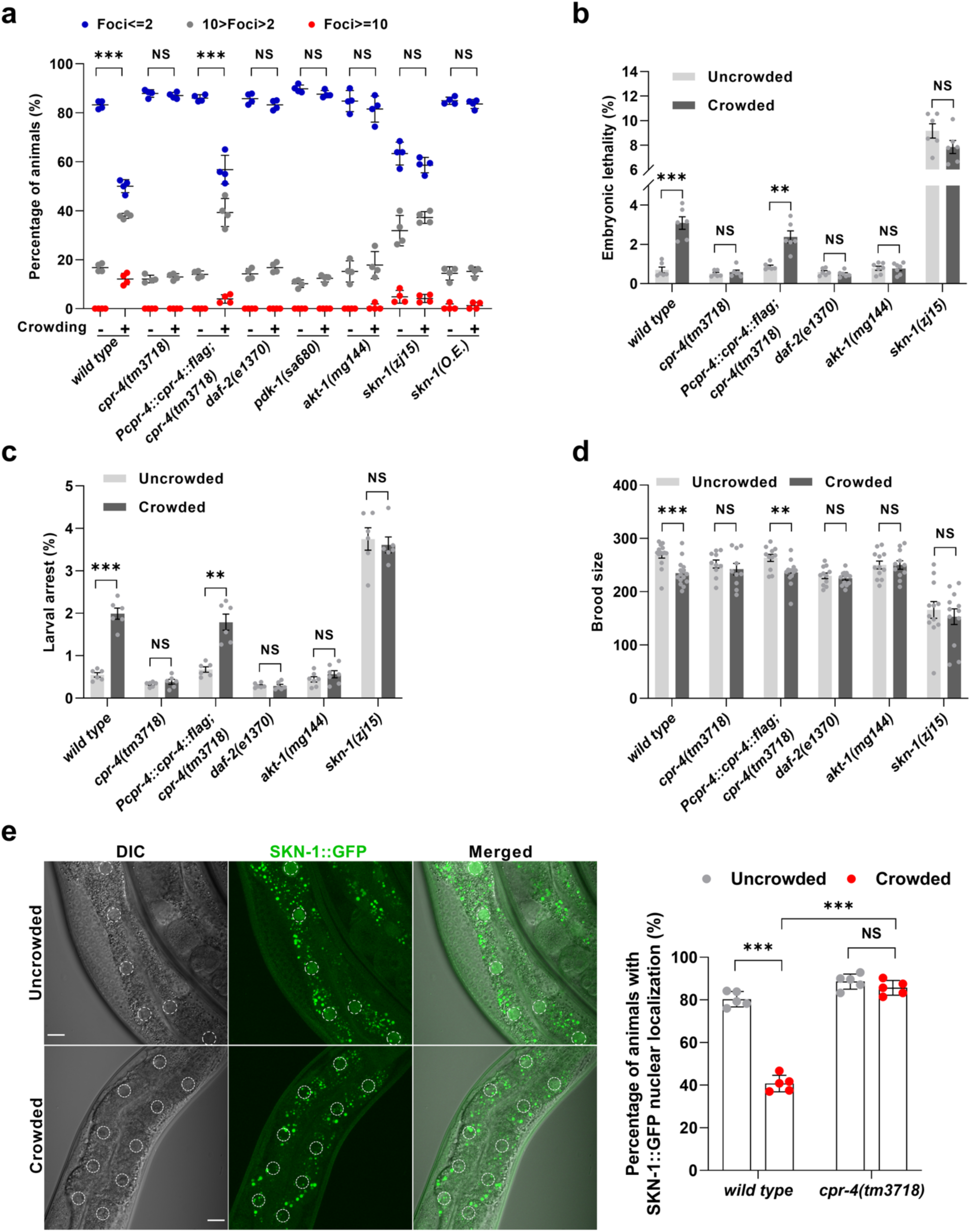
*cpr-4* and the *daf-2* insulin signaling pathway mediate the crowding responses. **a**, Percentages of *hus-1::neongreen* adult animals with the indicated genotype showing the number of gonad NeonGreen foci ≤2 (blue), between 3-9 (grey), and ≥10 (red), respectively. Animals were grown in the uncrowded and crowded conditions as indicated (each condition with four biologically independent repeats). *skn-1*(O.E.) indicates *skn-1* overexpression through an integrated transgene. **b**-**d**, Percentages of embryonic lethality (**b**) and larval arrest (**c**) observed in progeny and the brood sizes (**d**) of adult animals with the indicated genotype, which were grown in the uncrowded and crowded conditions, respectively. Six biologically independent experiments in **b** and **c**. **e**, Representative differential interference contrast (DIC), GFP, and merged images of wild-type animals expressing SKN-1::GFP grown in uncrowded and crowded conditions (left panel) and the percentages of animals with SKN-1::GFP nuclear localization in wild type and *cpr-4(tm3718)* animals grown in uncrowded and crowded conditions as indicated (right panel). Five biologically independent experiments at each condition were performed (n≥19 animals in each independent experiment). All animals carried the *smIs564 (*P*skn-1::skn-1::gfp)* transgene. Scale bars, 100 μm. The total numbers of adult animals analyzed from left to right are 119, 172, 117, 163, 152, 155, 144, 174, 156, 161, 158, 157, 145, 145, 159, and 158 in **a**, 120, 120, 120, 120, 120, 120, 120, 120, 140, 140, 120, and 120 in **b** and **c**, 14, 15, 10, 10, 11, 13, 11, 13, 12, 11, 12, and 12 in **d**, and 127, 127, 142, and 134 in **e**. Total numbers of embryos scored in **b**: 4064, 3758, 4875, 3680, 4345, 4572, 5716, 5179, 4749, 5021, 3618, and 3713 from left to right. Total numbers of larvae scored in **c**: 4038, 3653, 4866, 3672, 4336, 4545, 5700, 5174, 4711, 4981, 3367, and 3549 from left to right. Data are mean ± s.d. (**a**, **d**, **e)** or mean ± s.e.m (**b**, **c**). NS, not significant, \*\**P* < 0.01, \*\*\**P* < 0.001, two-sided *z*-test in **a** and two-sided, unpaired *t*-test in **b**, **c**, **d**, and **e**.

We examined the possibility that crowding might lead to hypoxia, which is known to induce DNA damage in some cellular contexts^34^. Using a hypoxia-responsive reporter, P*nhr-57*::GFP^35^, to detect the hypoxic levels in *C. elegans* animals, we found that the fluorescence intensity of P*nhr-57::GFP* in crowded animals was comparable to that in uncrowded animals or animals under the nonhypoxic condition (Fig. S1e, f), in contrast to the significantly elevated fluorescence intensity of P*nhr-57*::GFP in animals under the hypoxia condition (Fig. S1f). Moreover, real-time quantitative reverse transcription polymerase chain reaction (qRT-PCR) analysis to measure the transcription levels of key hypoxia-related genes, *hif-1, rhy-1, egl-9,* and *cysl-1*, showed that the transcription levels of these four genes were not significantly altered in animals in both crowded and uncrowded conditions (Fig. S1g). By contrast, the transcription of *cysl-1* and *hif-1* was upregulated and that of *egl-9* and *rhy-1* was downregulated, as expected, in animals under the hypoxia condition, compared with animals in the nonhypoxic condition (Fig. S1h). These results indicate that crowding stress does not lead to hypoxia and hypoxia stress response.

### DAF-22 is important for crowding-induced CPR-4 secretion and germ cell DNA damage

It has been shown that *C. elegans* animals can sense high population density and food availability to secrete multiple small molecule pheromones, which trigger different developmental changes, including dauer arrest when food supply is low^36–39^ or accelerated development when food supply is plentiful and animal population density is higher^40–42^. One of the genes crucial for mediating this density/food response, *daf-22,* encodes an ortholog of the human sterol carrier protein 2 (SCP2) and catalyzes peroxisomal fatty acid β-oxidation in the biosynthesis of pheromones^38,39,43^. We therefore investigated whether *daf-22* plays a role in regulating CPR-4 secretion in response to the crowding response. In crowded animals carrying a *daf-22(m130)* loss-of-function mutation, CPR-4 secretion was detectable but substantially lower than that in crowded wild type animals (Fig. S1i), and was not detected in uncrowded *daf-22(m130)* animals. These results suggest that loss of *daf-22* reduces CPR-4 secretion in response to the crowding stress. Consistent with this finding, we observed a slight increase of germ cell DNA damage in crowded *daf-22(m130)* animals compared with uncrowded *daf-22(m130)* animals or wild-type animals (Fig. 1d), indicating that *daf-22* is important for germ cell DNA damage induced by the crowding stress. Interestingly, uncrowded *daf-22(m130)* animals treated with C-CM from wild-type animals showed significantly increased germ cell DNA damage compared with that in uncrowded *daf-22(m130)* animals treated with NC-CM from wild-type animals and similar to the level of germ cell DNA damage observed in uncrowded wild-type animals treated with C-CM (Fig. 1d). These results together suggest that *daf-22* acts upstream of or in parallel to *cpr-4* to regulate CPR-4 secretion and germ cell DNA damage in response to the crowding stress.

### The insulin signaling pathway is required for crowding-induced reduction in animal survival and fecundity

We next examined the consequence of increased germline DNA damage and found that animals grown in the crowded condition exhibited increased embryonic lethality and larval arrest, and as reported previously^44^, decreased brood sizes, compared with those grown in the uncrowded condition (Fig. 2b-d). We rescued animals with varying degrees of germline DNA damage in both uncrowded and crowded conditions after microscopy screens and found that the severity of germ cell DNA damage positively correlated with the severity of embryonic lethality, larval arrest and brood size reduction (Fig. 1e-g; Fig. S1j-l), with crowded animals having 10 or more HUS-1::NeonGreen foci showing the highest embryonic lethality (40.3%), larval arrest (8.3%) and brood size reduction (an average of 53 progeny). In comparison, uncrowded and crowded animals with two or less HUS-1::NeonGreen foci had 2.3% and 2.6% embryonic lethality, 2.0% and 2.1% larval arrest, and an average of 267 and 250 progeny, respectively (Fig. 1e-g; Fig. S1j-l). Similarly, the *cpr-4(tm3718)* mutation suppressed these crowding-induced developmental defects (Fig. 2b-d), which were restored by the P*cpr-4::cpr-4::flag* transgene. Therefore, increased CPR-4 secretion and germ cell DNA damage induced by crowding exert an adverse effect on animal development and reproduction.

The insulin/IGF-1 signaling pathway (IIS pathway) is involved in regulating multiple important cellular processes, including stress response, metabolism and life span, and is conserved from nematodes to humans^45–48^. The *C. elegans* homologue of human insulin/IGF-1 receptor, DAF-2, is a key component of the insulin/IGF-1 signaling pathway^49^. Reduced DAF-2 activity has been shown to increase stress resistance, extend lifespan, and block radiation-induced bystander effects^31,45,48^. We found that crowding-induced germ cell DNA damage, embryonic lethality, larval arrest, and brood size reduction were all strongly suppressed by a *daf-2(e1370)* loss-of-function mutation (Fig. 2a-d), indicating that DAF-2 is required for the crowding responses. Moreover, a loss-of-function mutation in *pdk-1 (sa680)* and a gain-of-function mutation in *akt-1 (mg144),* two downstream components of the DAF-2 IIS pathway^50,51^, also strongly suppressed the crowding-induced DNA damage and developmental defects (Fig. 2a-d and Fig. S2b-d). These results indicate that the DAF-2 insulin/IGF-1 signaling pathway is crucial for mediating the crowding responses. Because *daf-2(e1370)* and *akt-1 (mg144)* also blocked germ cell DNA damage induced by secreted CPR-4 in C-CM (Fig, 1d; Fig. S2e), these results suggest that the *daf-2* insulin signaling pathway acts downstream of or in parallel to *cpr-4* to mediate crowding responses.

DAF-16 is one of the major downstream transcription factors in the DAF-2 IIS pathway that regulates multiple important cellular processes^52,53^. Interestingly, a strong loss-of-function mutation in *daf-16 (mu86)* did not block or enhance crowding-induced DNA damage and developmental defects (Fig. S2a-d), indicating that DAF-16 is dispensable for the crowding responses. In the *daf-16(mu86); daf-2(e1370)* double mutant, crowding-induced germ cell DNA damage remained strongly suppressed (Fig. S2a), confirming that the crowding-induced DNA damage is dependent on *daf-2* but independent of *daf-16*. We then examined another transcription factor, SKN-1, that also acts downstream of the DAF-2 signaling pathway to regulate stress responses and longevity^54–56^. In a *skn-1(zj15)* partial loss-of-function mutant^57^, a higher percentage of animals showed increased germ cell DNA damage under the noncrowded condition, which was not exacerbated by the crowding stress (Fig. 2a). This result suggests that reduced *skn-1* activity alone is sufficient to cause germ cell DNA damage and that the crowding stress may induce DNA damage through inhibiting the *skn-1* activity. On the other hand, overexpression of *skn-1* blocked both crowding- and C-CM-induced germ cell DNA damage (Fig. 2a; Fig. S2e), indicating that SKN-1 acts downstream of or in parallel to the crowding signaling pathway to protect against germ cell DNA damage. The *skn-1(zj15)* mutation on its own caused increased embryonic lethality and larval arrest and decreased brood sizes in the uncrowded conditions, which were not exacerbated by the crowding stress (Fig. 2b-d). These results together suggest that *skn-1* is important for maintaining genome integrity and normal animal development and that these important *skn-1* functions are compromised during the crowding stress.

Since SKN-1 acts in the nuclei to regulate transcription of genes important for development, stress responses, and longevity^54–56^, we investigated nuclear localization of SKN-1 in animals carrying an integrated transgene expressing SKN-1::GFP (P*skn-1::skn-1::gfp)* under uncrowded and crowded conditions. Notably, the percentage of animals with intestinal nuclear SKN-1::GFP was significantly lower in crowded animals than that in uncrowded animals (Fig. 2e), suggesting that the crowding stress causes decrease of the SKN-1 nuclear localization. In contrast, in the *cpr-4* mutant, the percentages of animals with nuclear SKN-1::GFP were comparable in both crowded and uncrowded conditions, indicating that loss of *cpr-4* negates crowding-induced decrease of SKN-1 nuclear localization and that crowding inhibits SKN-1 activity through CPR-4. Consistent with the finding that *cpr-4* modulates the SKN-1 activity in response to the crowding stress, the fluorescence intensity of the P*_gst-4_*GFP transcriptional fusion, widely used as an indicator of the SKN-1 transcriptional activity^58,59^, decreased significantly in crowded animals, compared with that in uncrowded animals, but was not reduced in *cpr-4* mutant animals in both crowded and uncrowded conditions (Fig. S2f), confirming that the crowding stress leads to downregulation of the SKN-1 activity through *cpr-4*. Taken together, these data suggest that the crowding stress and CPR-4 induce germ cell DNA damage and related developmental deficiencies through impairing the protective functions of SKN-1.

### Crowding and CPR-4 increase genome mutation rates

Recognizing that DNA damage in germ cells can generate genetic mutations and increase genome mutation rates, we investigated whether crowding stress and increased secretion of CPR-4 caused a rise in genome mutations. We grew progeny of single L4 stage wild-type (N2) and *cpr-4(tm3718)* animals in both crowded (C) and uncrowded (NC) conditions continuously for multiple generations (Fig. 3a). At the 10^th^ generation, multiple N2 and *cpr-4(tm3718)* animals from each condition were randomly selected and grown clonally. Their progeny were subjected to whole-genome sequencing analyses to ascertain accumulated genome mutations. For comparison, first-generation N2 (N2^F1^) and *cpr-4* mutant (*cpr-4*^F1^) animals grown in uncrowded conditions were also sequenced.

**Fig. 3.**
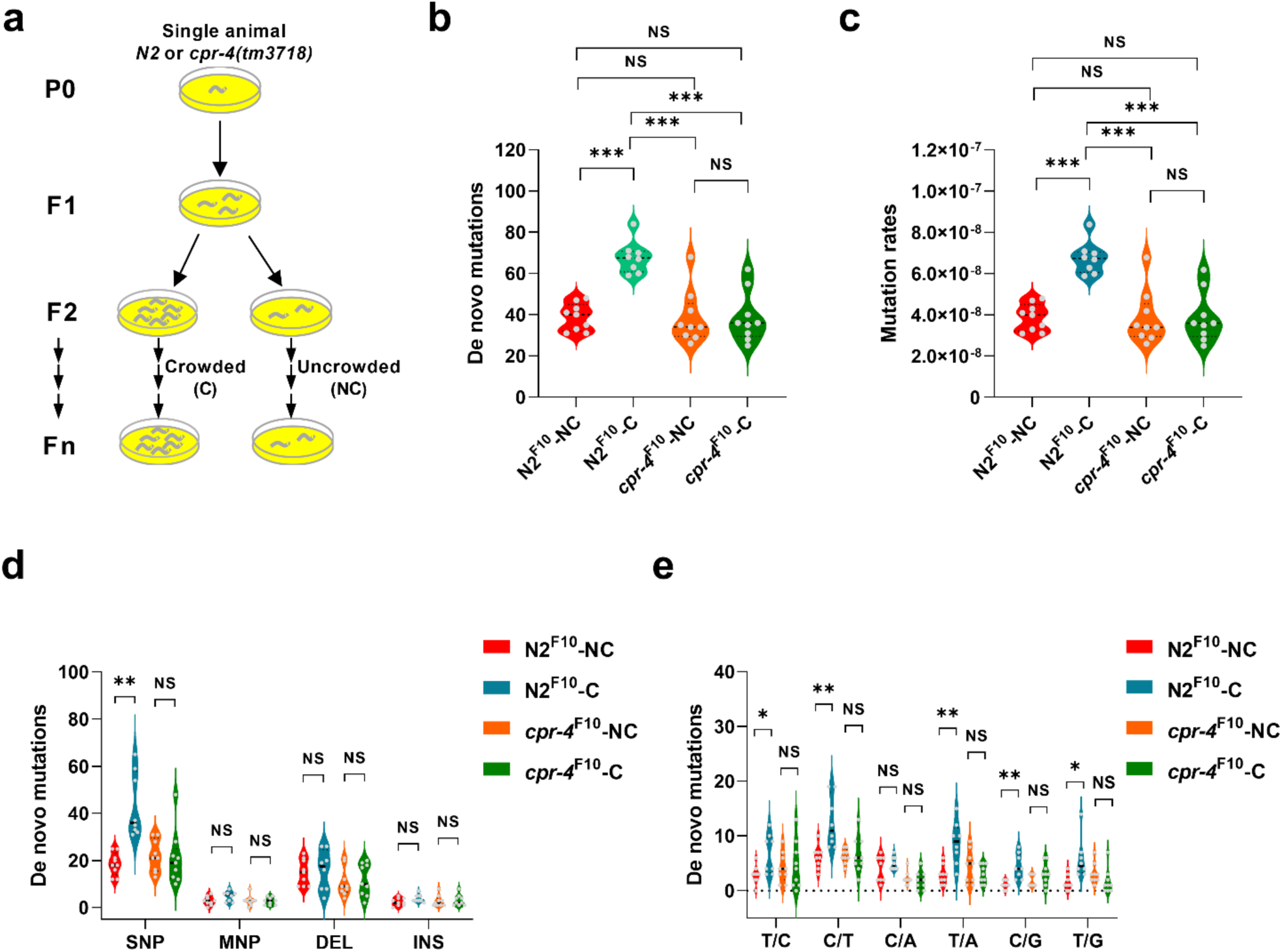
*cpr-4* promotes generation of de novo genome mutations and increases mutation rates under persistent crowding stress. **a**, A schematic diagram showing the continuous growth of wild-type (N2) and *cpr-4(tm3718)* animals in uncrowded and crowded conditions, respectively, for multiple generations. **b**, **c**, The number of de novo mutations (**b**) and the mutation rate per base pair (bp) (**c**) identified in N2 and *cpr-4(tm3718)* animals at the 10^th^ generation of continuous growth in crowded (C) and uncrowded (NC) conditions. **d**, **e**, The numbers of four major types of de novo mutations, single nucleotide polymorphism variants (SNP), multiple nucleotide polymorphism variants (MNP), deletions (DEL), and insertions (INS) (**d**), and six different types of SNP changes (**e**) identified in N2 and *cpr-4(tm3718)* animals at the 10^th^ generation of continuous growth in crowded and uncrowded conditions. Animals from each genotype and growing condition were randomly chosen and whole-genome sequenced and analyzed (**b**-**e**). n=9 (N2^F10^-NC), 8 (N2^F10^-C), 9 (*cpr-4*^F10^-NC), and 9 (*cpr-4*^F10^-C) from two biologically independent experiments. NS, not significant, \**P* < 0.05, \*\**P* < 0.01, \*\*\**P* < 0.001, two-sided, unpaired *t*-test.

In *C. elegans*, animals acquire approximately 0.8 to 2 spontaneous germline mutations per generation^60–63^. To determine whether crowding induces more genome mutations and increases mutation rates, we analyzed the numbers of de novo mutations in N2 and *cpr-4* mutant animals grown in uncrowded and crowded conditions for ten generations. In uncrowded N2^F10^-NC animals, we detected an average of 38.8 de novo mutations, which corresponds to a mutation rate of approximately 3.9 ×10^-8^ per site per generation, in line with the spontaneous mutation rates (0.8×10^-8^ to 2.0×10^-8^) observed previously^60–63^ (Fig. 3b, c). Notably, significantly more de novo mutations were found in crowded N2^F10^-C animals (an average of 67.8) than in uncrowded N2^F10^-NC animals, with an average increase of 29 germline mutations per animal and 75% increase in mutation rates per site per generation (Fig. 3b, c). These findings suggest that crowding stress promotes generation of more genome mutations and increase of mutation rates. In contrast, the numbers of de novo mutations and the mutation rates per site per generation detected in uncrowded *cpr-4(tm3718)*^F10^-NC animals and crowded *cpr-4(tm3718)*^F10^-C animals showed no significant difference (Fig. 3b, c), indicating that loss of *cpr-4* blocks generation of crowding-induced de novo mutations and elevated mutation rates. Importantly, the numbers of de novo mutations generated and the mutation rates seen in *cpr-4(tm3718)*^F10^-C animals are comparable to those observed in N2^F10^-NC animals, but significantly lower than those found in N2^F10^-C animals (Fig. 3b, c), confirming the crucial role of *cpr-4* in mediating the surge of crowding-induced de novo mutations and mutation rates.

We investigated the nature of genome mutations induced by crowding stress and CPR-4 and found that single nucleotide polymorphism variants (SNP), were significantly higher in N2^F10^-C animals than in N2^F10^-NC animals (Fig. 3d), whereas the numbers of deletions (DEL) and insertion mutations (INS) and multiple nucleotide polymorphism variants (MNP) found in N2^F10^-C and N2^F10^-NC animals were comparable (Fig. 3d). Consistent with the finding that CPR-4 plays an important role in mediating increase of genome mutations induced by crowding stress, we did not observe significant difference in generation of these four types of de novo mutations among *cpr-4*^F10^-NC, *cpr-4*^F10^-C, and N2^F10^-NC animals (Fig. 3d). These results suggest that crowding stress and CPR-4 induce increased generation of SNPs in the genomes.

We further analyzed the types of SNP variants induced by crowding stress through examining T to A (T/A or A/T in reverse), T to C (T/C or A/G in reverse), T to G (T/G or A/C in reverse), C to T (C/T or G/A in reverse), C to A (C/A or G/T in reverse), and C to G (C/G or G/C in reverse) changes. We found that except C to A (C/A or G/T) changes, the numbers of all other types of SNPs were significantly higher in crowded N2^F10^-C animals than in uncrowded N2^F10^-NC animals, with the C to T (or G to A) changes showing the largest increase. As expected, we observed no significant difference in the numbers of six types of SNP mutations produced among *cpr-4*^F10^-NC, *cpr-4*^F10^-C, and N2^F10^-NC animals (Fig. 3e). Taken together, these results provide strong evidence that CPR-4 mediates crowding-induced increase of genome mutations, leading to higher mutation rates in animals grown under persistent crowding stress.

### Crowding-induced de novo mutations exhibit biased distribution in the genome

We investigated the distribution pattern of crowding-induced de novo mutations in the genome by comparing the occurrences of de novo mutations in gene bodies and intergenic regions. We found that in crowded N2^F10^-C animals the numbers of de novo mutations in the intergenic regions were significantly higher than those in uncrowded N2^F10^-NC animals (Fig. 4a). In contrast, the numbers of de novo mutations in gene bodies were comparable between N2^F10^-NC and N2^F10^-C animals (Fig. 4a). Compared with uncrowded N2^F10^-NC animals, the percentage of de novo mutations in crowded N2^F10^-C animals was significantly higher in the intergenic regions but lower in the gene bodies (Fig. S3a), indicating that persistent crowding stress induces biased mutation distribution towards the intergenic regions in crowded animals.

**Fig. 4.**
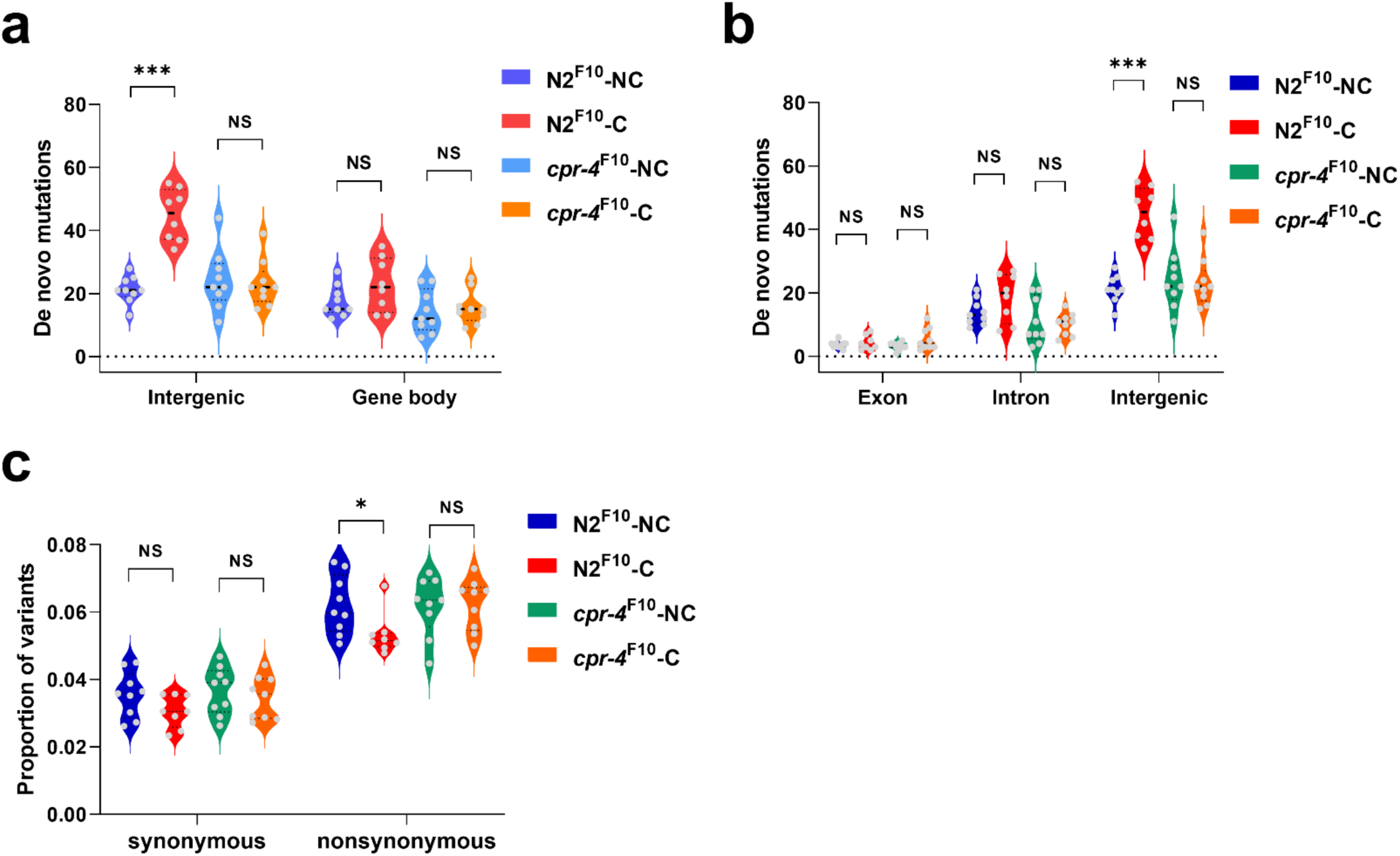
Persistent crowding stress and *cpr-4* drive biased mutation distributions in the genomes. **a**, **b**, Numbers of de novo mutations identified in the intergenic regions and gene bodies (**a**) or in exons, introns, and intergenic regions (**b**) in N2 and *cpr-4(tm3718)* animals at the 10th generation of continuous growth in crowded (C) and uncrowded (NC) conditions, respectively. **c**, Proportions of synonymous and nonsynonymous variants identified in N2 and *cpr-4(tm3718)* animals at the 10th generation of continuous growth in uncrowded and crowded conditions. n=9 (N2^F10^-NC), 8 (N2^F10^-C), 9 (*cpr-4*^F10^-NC), and 9 (*cpr-4*^F10^-C) from two biologically independent experiments. NS, not significant, \**P* < 0.05, \*\*\**P* < 0.001, two-sided, unpaired *t*-test (**a**, **b**) or Wilcoxon rank-sum test (**c**).

In comparison, 10 generations of continual growth of *cpr-4(tm3718)* animals in both uncrowded (*cpr-4*^F10^-NC) and crowded (*cpr-4*^F10^-C) conditions produced similar numbers of de novo mutations in both intergenic regions and gene bodies (Fig. 4a), suggesting that inactivation of *cpr-4* eliminates the biased mutation distribution induced by persistent crowding stress. Similarly, the percentages of de novo mutation in the intergenic regions and gene bodies show no significant difference between *cpr-4*^F10^-NC and *cpr-4*^F10^-C animals (Fig. S3a). These results indicate that genome mutations induced by CPR-4 exhibit biased distribution toward the intergenic regions under persistent crowding stress, which may select against mutations in coding regions that compromise animal development, growth and fitness and lead to their reduced distribution in gene bodies.

### Crowding-induced de novo mutations are subject to crowding selection

To further investigate this biased mutation distribution, we analyzed the distribution of crowding-induced mutations within gene bodies and found that N2^F10^-C animals had significantly lower percentages of de novo mutations in introns, but significantly higher percentages of de novo mutations in the intergenic regions than N2^F10^-NC animals (Fig. S3b), indicating that persistent crowding stress decreases distribution of de novo mutations in gene bodies. Interestingly, N2^F10^-C animals had similar numbers of de novo mutations in exons and introns like N2^F10^-NC animals, but had significantly more de novo mutations in the intergenic regions than N2^F10^-NC animals (Fig. 4b). These results suggest that persistent crowding stress favors generation and accumulation of genome mutations in the intergenic regions, but disfavors mutations in exons and introns, potentially due to negative crowding selection. Since mutations in exons, and to a lesser extent, in introns often impair or compromise the functions of proteins or RNAs crucial for animals’ development, growth, and survival, they are more likely to be eliminated under persistent crowding selection. Consistent with this notion, the proportion of nonsynonymous variants in N2^F10^-C animals was significantly lower than that in N2^F10^-NC animals, whereas the proportions of synonymous variants were similar in N2^F10^-C and N2^F10^-NC animals (Fig. 4c). These results suggest that mutations in the gene bodies of N2^F10^-C animals experienced stronger negative selection during 10-generation persistent crowding stress. In comparison, *cpr-4*^F10^-C animals showed comparable percentages of de novo mutations and numbers of de novo mutations in gene bodies and intergenic regions, as well as similar proportions of both nonsynonymous and synonymous variants to those of *cpr-4*^F10^-NC animals (Fig. 4b,c and Fig. S3b), indicating that *cpr-4* is crucial for the crowding selection that leads to increased distribution of genome mutations in the intergenic regions and their decreased presence in gene bodies under persistent crowding stress.

To test the hypothesis that crowding selection leads to decreased presence of de novo mutations in gene bodies that impair growth, survival, and other crucial functions of animals, we grew N2 and *cpr-4*(*tm3718)* animals in both crowded and uncrowded conditions continuously for 15 and 30 generations, respectively, and screened animals for visible phenotypes, which potentially have mutations in gene bodies. We observed significantly higher percentages (9% at the 15^th^ generation and 12% at the 30^th^ generation) of N2 animals with visible phenotypes under the crowded condition than N2 animals grown in the uncrowded condition (1.7% at both 15^th^ and 30^th^ generations; Fig. 5a). In contrast, the percentages of abnormal animals observed in the *cpr-4(tm3718)* mutant were low in both crowded and noncrowded conditions (1.2%-1.8%), comparable to those of N2 animals grown in the uncrowded condition (Fig. 5a). These results suggest that under the crowded condition CPR-4-induced mutagenesis yields significantly higher numbers of animals with visible phenotypes. Numerous different phenotypes were observed, including increased frequency of males (4.5-5.2% in the crowded condition versus 0.5-0.6% in the uncrowded condition), animals that were uncoordinated in locomotion or immobile, animals that had blisters or multiple vulvas, and animals that were small or dumpy (Fig. S4a, b), all of which are phenotypes typically observed in mutants derived from chemical mutagenesis^64^. Importantly, approximately 3.3% of those visible phenotypes from N2 animals grown under the crowded condition were heritable (3 out of 90 selected for further analysis at the 15^th^ generation and 4 out of 117 at the 30^th^ generation; Fig. 5a and Fig. S4c), whereas no heritable mutation was found from abnormal animals at the 15^th^ generation (17 animals) and the 30^th^ generation (17 animals) grown under the uncrowded condition. There was no heritable mutation found in abnormal *cpr-4(tm3718)* animals at the 15^th^ generation (12 from the noncrowded condition and 18 from the crowded condition) and the 30^th^ generation (14 from the noncrowded condition and 16 from the crowded condition), suggesting that CPR-4 is crucial for generating heritable mutations in animals grown under persistent crowding stress.

**Fig. 5.**
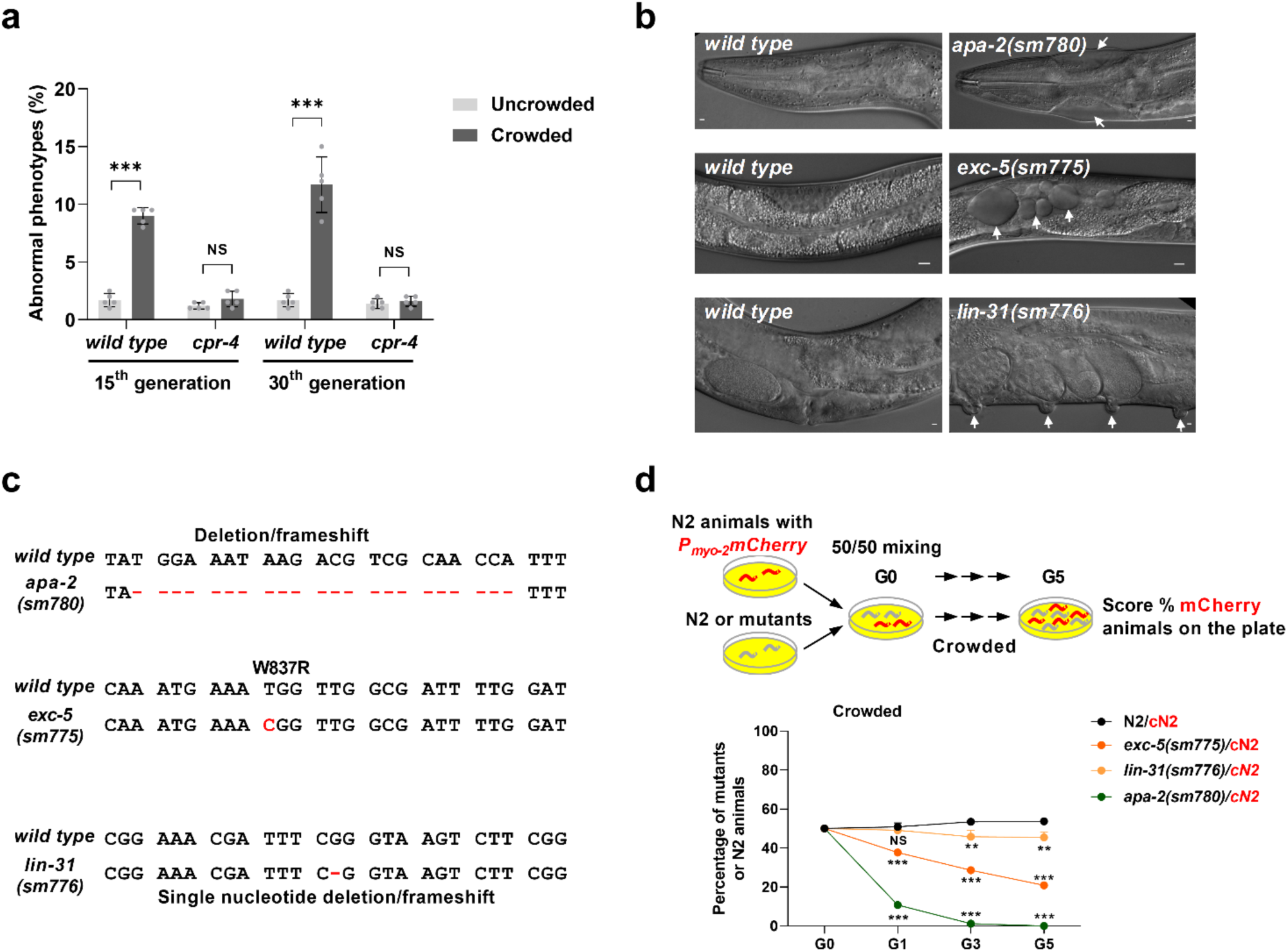
Negative crowding selection against de novo mutations in gene bodies induced by persistent crowding stress and *cpr-4*. **a**, Percentages of N2 and *cpr-4(tm3718)* animals with abnormal phenotypes identified at the 15^th^ and the 30^th^ generations of continuous growth in crowded and uncrowded conditions, respectively. Five biologically independent experiments were performed in each condition (n=1000 animals scored). **b**, Representative differential interference contrast images showing mutant phenotypes (indicated by arrows, right panels) caused by heritable mutations in three different genes generated under continuous crowded conditions. Corresponding images of wild-type animals are shown in left panels. Scale bars, 20 μm. **c**, Molecular lesions of three genetic mutations shown in (**b**), confirmed by DNA sequencing, as compared with the corresponding wild-type DNA sequences. **d**, Five-generation growth competition assays between two groups of animals grown in crowded conditions, in which one group of 50 animals were labeled by mCherry expressed from a single copy P*_myo-2_*mCherry integrated transgene (cN2 in red) and the other group of 50 animals were not labeled (N2 or mutant animals). The percentages of unlabeled animals in the populations at the first (G1), third (G3), and fifth (G5) generations were scored (*n* = 300 in each experiment). The heritable genetic mutations, *exc-5(sm775)*, *lin-31(sm776)*, and *apa-2(sm780)*, identified from the 15^th^ and 30^th^ generations were used in the growth competition assays. Three biologically independent experiments were performed for each pair of animals and condition. Data are mean ± s.d. (**a**, **d**). NS, not significant, \*\**P* < 0.01, \*\*\**P* < 0.001, two-sided, unpaired *t*-test (**a)** or binomial generalized linear model (logit) followed by the Dunnett-type comparisons test in **d**.

To characterize these heritable mutations, we performed whole genome sequencing of all seven mutants (Fig. S4c) and genetic mapping of three of them (*sm775, sm776*, and *sm780*) to determine their approximate chromosomal positions. A 22 base-pair deletion in the *apa-2* gene, a frameshift mutation, was found in the *sm780* mutant that exhibited a phenotype of blistered cuticle at its head (Fig. 5b,c) and was mapped to chromosome X^65^. This defect was rescued by a fosmid containing the *apa-2* gene (Fig. S4d), indicating that the 22 bp deletion in *apa-2* is responsible for the phenotype. In the *sm775* mutant that displayed a phenotype of multiple vacuoles and was mapped to chromosome IV, a T to C transition that results in Trp 837 to Arg substitution was identified in the *exc-5* gene (Fig. 5b, c), which has the same mutant phenotype^66^. A fosmid containing the *exc-5* gene can fully rescue this vacuolar defect (Fig. S4e). In the *sm776* mutant that had a multivulva defect and was mapped to chromosome II, a single nucleotide deletion, resulting in a frameshift, was found in the coding region of the *lin-31* gene (Fig. 5b,c), which is known to have a multivulva mutant phenotype^67^. We confirmed that *sm776* is an allele of *lin-31* through a complementation test with the canonical *lin-31(n301)* mutation. The phenotypes of the remaining heritable mutants were listed in Fig. S4c. Together, these results suggest that the crowding stress and CPR-4 increase generation of heritable genome mutations, including those affecting the exons of genes.

We then examined whether the three heritable mutations in the coding regions affected animal growth using a five-generation growth competition assay. In this assay, N2 animals were labeled with mCherry at the head (named cN2 as cherry-labeled N2 animals) using a single-copy mCherry reporter (P*_myo-2_*mCherry) inserted into the *C. elegans* genome through CRISPR/Cas9 genome editing^68^ and placed with unlabeled N2 animals at 1:1 ratio in the crowded condition for 5 consecutive generations. The cN2 animals were indistinguishable in growth rate from unlabeled N2 animals in the crowded condition (black line, Fig. 5d), each comprising approximately 50% of the population in alternate generations examined (G1, G3, and G5 as generation 1, 3, and 5, respectively). Importantly, all three mutants displayed less competitive growth in the crowded conditions compared with cN2 animals (Fig. 5d), with the *apa-2(sm780)* mutant showing the severest growth defect. Only 1% of *apa-2(sm780)* animals were present in the mixed population at the 3^rd^ generation and cN2 animals completely took over the population by the 5^th^ generation (the dark green line, Fig. 5d). These results demonstrate that under persistent crowding stress animals with harmful or unfavorable mutations in coding regions, which impair animal development, growth and functions, will face strong negative selection that leads to their dwindling presence or elimination from the population. After multiple generations of high-density selection, this can result in less mutations detected in gene bodies and more mutations found in the intergenic regions, compared with their uncrowded cohorts (Fig. 4 and Fig. S3). Such biased mutation distribution across the genome under persistent crowding stress underlies the observed genome evolution to generate animals that grow competitively in the crowded environment.

### Animals under persistent crowding stress acquire inducible growth advantage

To further analyze if animals under persistent crowding stress evolve to grow competitively in the crowded environment, we continuously grew both N2 and *cpr-4 (tm3718)* animals with or without the single-copy P*_myo-2_*mCherry transgene in crowded and uncrowded conditions for 30 generations, respectively, and then examined which population of animals became more adapted to the crowded condition and displayed a growth advantage using the five-generation growth competition assay discussed above (Fig. 5d and Fig. 6a). The growth rate of first generation mCherry-labeled cN2 animals (cN2^F1^) were indistinguishable from that of unlabeled N2 animals (N2^F1^) in both crowded and uncrowded conditions for 5 consecutive generations (black lines, Fig. 6b, c), each comprising 50% of the population in alternate generations examined. After 30 generations of evolution in both crowded and uncrowded conditions, cN2 animals grown in the crowded condition (cN2^F30^-C) clearly had a growth advantage over N2 animals grown in the uncrowded condition (N2^F30^-NC), outcompeting N2^F30^-NC animals by 22% by the 5^th^ generation in the growth competition assay (orange line, Fig. 6b). A similar level of growth advantage by N2^F30^-C animals over cN2^F30^-NC animals was observed (blue line, Fig. 6b). Interestingly, when the same two pairs of animals (N2^F30^-NC/cN2^F30^-C and N2^F30^-C/cN2^F30^-NC) were placed in the uncrowded condition, they showed comparable growth rates, each taking up approximately 50% of the population (Fig. 6c). These results indicate that the growth advantage observed in cN2^F30^-C and N2^F30^-C animals is not permanent and is inducible by the crowding stress, which enables them to rapidly adapt to the crowded environment. As expected, when the noncrowded pair (N2^F30^-NC /cN2^F30^-NC) or the crowded pair (N2^F30^-C/cN2^F30^-C) were placed in the crowded and uncrowded conditions, respectively, they showed similar growth rates (Fig. S5a, b).

**Fig. 6.**
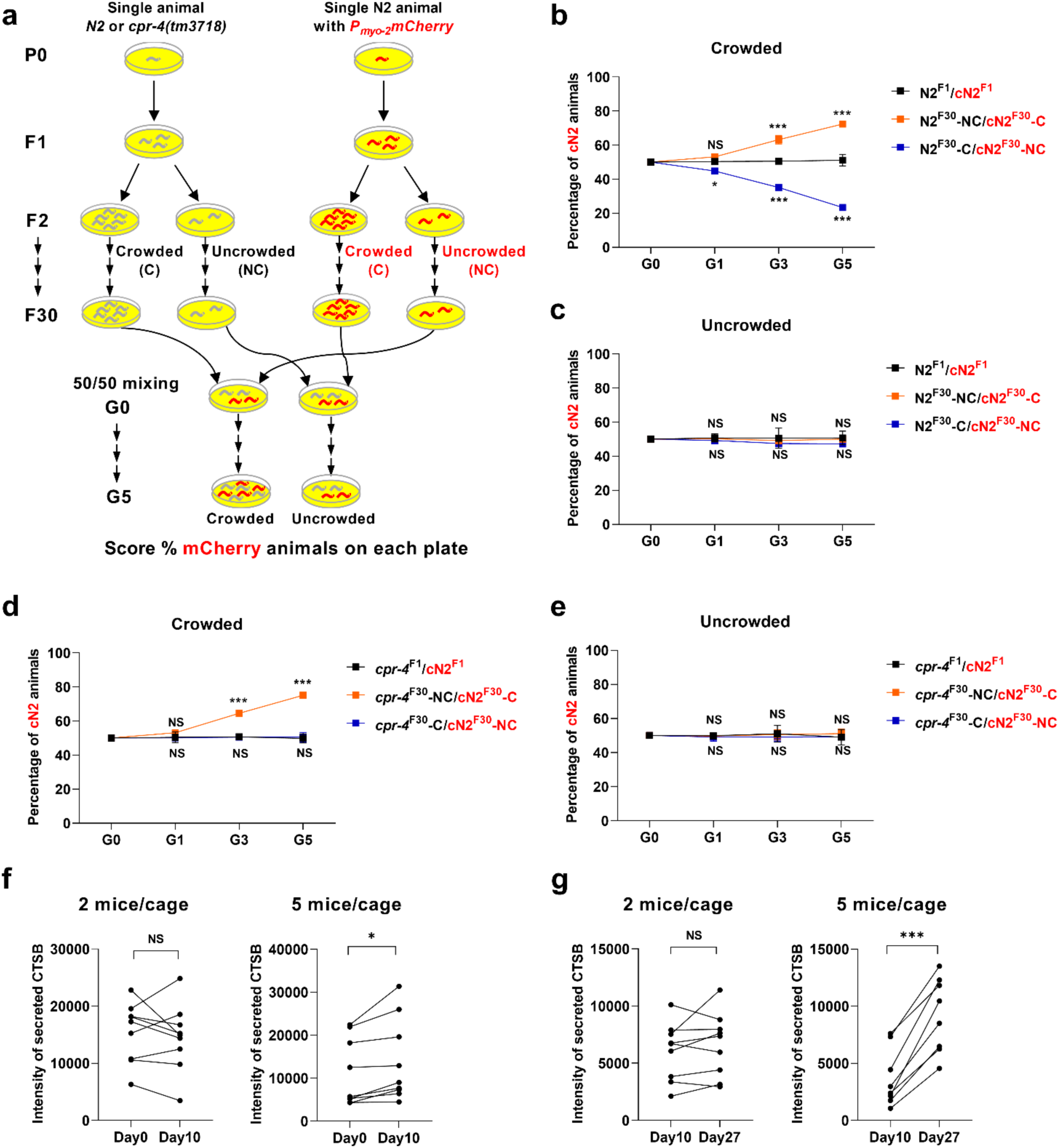
*cpr-4* promotes inducible growth advantage after 30 generations of continuous crowding stress. **a**, A schematic diagram of N2 and *cpr-4(tm3718)* animals with or without the single-copy P*_myo-2_*mCherry integrated transgene grown in crowded (C) and uncrowded (NC) conditions continuously for 30 generations, respectively. Subsequently, two selected groups of animals (initially with 50 animals each) were subjected to five-generation growth competition assays as in Fig. 5d. **b**-**e**, Five-generation growth competition assays between the indicated groups of animals in crowded (**b**, **d**) and uncrowded (**c**, **e**) conditions. in which one group of animals were labeled by mCherry (cN2 in red). The percentages of mCherry-labeled animals in the populations at the first (G1), third (G3), and fifth (G5) generations were scored (*n* = 300 in each experiment). First generation (F1) animals were used as unconditioned controls. Animals at the 30^th^ generation (F30) grown in either crowded (C) or uncrowded (NC) conditions were used in the growth competition assays. Three biologically independent experiments were performed for each pair of animals and condition. **f**, **g**, Increased secretion of cathepsin B in mice under crowding stress. Plasma of mice housed in 2 mice per cage and 5 mice per cage were collected on Day 0 (before crowd conditioning), Day 10, and Day 27. The levels of secreted CTSB in mouse plasma were detected by immunoblotting (IB) using a monoclonal antibody to human CTSB (see Fig. S5e, f). 8-9 mice were analyzed for each group. The intensities of the secreted CTSB bands from the same mouse at three different time points were quantified and compared in pairs. Data are mean ± s.d. (**b**, **c**, **d**, and **e**). NS, not significant, \**P* < 0.05, \*\*\**P* < 0.001, binomial generalized linear model (logit) followed by the Dunnett-type comparisons test (**b**, **c**, **d**, and **e**) or two-sided, paired *t*-test (**f**, **g)**.

On the other hand, *cpr-4(tm3718)* animals that went through 30 generations in the crowded condition (*cpr-4*^F30^-C) showed no growth advantage over cN2^F30^-NC animals in both crowded and uncrowded conditions (blue lines, Fig. 6d, e) and were quickly outgrown by cN2^F30^-C animals in the crowded environment (green line, Fig. S5d). These results indicate that loss of *cpr-4* abolishes crowding-inducible growth advantage seen in N2^F30^-C animals. Moreover, *cpr-4(tm3718)* animals grown in the uncrowded condition for 30 generations (*cpr-4*^F30^-NC) had a comparable growth rate to that of cN2^F30^-C animals in the uncrowded condition (Fig. 6e), but were rapidly outgrown by cN2^F30^-C animals in the crowded environment (orange line, Fig. 6d), confirming that the growth advantage seen in crowd-conditioned N2 animals needs to be induced by the crowding stress. Together, these results suggest that animals under persistent crowding stress acquire an inducible growth advantage during evolution through a CPR-4-dependent mechanism to respond rapidly and competitively to the crowding stress.

### Increased secretion of cathepsin B in mice under crowding stress

Because the cathepsin B (CTSB) protease, the CPR-4 homologue, is highly conserved in the animal kingdom (Fig. S6; at least 45% sequence identity among cathepsin B proteins from different species), we examined whether secretion of CTSB is elevated in crowded conditions in other species, such as mammals. It has been reported that the expression levels of cathepsin B were significantly higher in the hippocampi of mice housed in a crowded condition with six mice per cage than those of mice living in a less crowded condition of two mice per cage^69^. Consistent with this finding, we observed significant increases of CTSB secretion in the blood of mice housed in the crowded condition of five mice per cage in two consecutive time spans of 10 days and 17 days (Fig. 6f, g and Fig. S5e, f). In contrast, no significant changes in CTSB secretion were detected in mice living in a less crowded condition of two mice per cage in the same time spans (Fig. 6f, g and Fig. S5e, f). Because in natural habitat mice enjoy an activity range of at least 1000 m^2^ from their home base^25^, whereas laboratory mice live in approximately 0.3 m^2^ restricted cage space, both cage conditions could be considered as crowded, with 5 mice per cage being the substantially more crowded one. These results demonstrate that mice also respond to the crowding stress by secreting CTSB, which could be a general crowding stress response in animals. Whether elevated circulating CTSB has a causal role in poorer reproductive outcomes shown to occur in mice when housed in crowded conditions^25,26^, and thus might represent a conserved mechanism, remains to be determined.

## Discussion

The regulation of population density is crucial for the reproduction, survival, and fitness of different species, as the population of a species cannot increase indefinitely. Density-dependent regulation has been a fundamental principle in modulating the population dynamics in the animal kingdom^14-17^, but the underlying molecular mechanisms have remained elusive^14,15^.

In this study, we show that when the population density of the nematode *C. elegans* reached a certain threshold, animals started to have genomic DNA damage in their germ cells, which increased in a population density-dependent manner. The genomic DNA damage in germ cells was followed by increased embryonic lethality and larval arrest and decreased brood size in the next generation, which are the most common deficiencies seen in a population with high density and could alleviate the impact of overpopulation^9-13^. Moreover, the severity of these deficiencies positively correlated with the degree of germ cell DNA damage. Importantly, these crucial events also correlated with the density-dependent secretion of the CPR-4 protein by late larval and adult animals with proliferating germ cells and reproductive capability. CPR-4 is the key mediator of radiation-induced bystander effects^31^ and a clastogenic factor that can promote chromosomal DNA breaks in germ cells non-cell-autonomously and across animals^32^. Inactivation of the *cpr-4* gene abolished all these density-dependent DNA damage and developmental deficiencies, indicating that CPR-4 is a crucial density-responding factor that induces genomic DNA damage in germ cells to promote density-dependent changes in crowded animals. Moreover, the DAF-2 insulin-like growth factor receptor signaling pathway and one of its downstream transcription factors, SKN-1, which are involved in mediating stress responses in *C. elegans*^45,48,54,56,70^, are required for CPR-4 to trigger these density-dependent DNA damage and developmental deficiencies. Therefore, our study identifies a density-responding molecule and a signaling pathway that regulates the responses of animals to the crowding stress.

Genetic variations are the driving force of evolution during natural selection^71^, but the molecular mechanisms underlying the generation of genetic variations are poorly characterized^72^. In this study, we show that CPR-4, a clastogenic factor, was secreted in response to the crowding stress to promote germ cell DNA damage, which significantly increased genome mutation rates and the number of *de novo* genetic mutations when the crowding stress persisted for multiple generations (Fig. 3). Loss of *cpr-4* prevented germ cell DNA damage, increase of genome mutation rates and *de novo* mutations, biased mutation distribution in the genome (Fig. 4 and Fig. S3), and inducible adaptive responses of animals to the crowding stress (Fig. 6 and Fig. S5), revealing a pivotal role of *cpr-4* in promoting generation of genetic variations, genome evolution, and adaptive changes. Indeed, our whole genome sequencing analyses of animals from 10 generations of continuous growth in both crowded and uncrowded conditions show that CPR-4-induced DNA damage and mutagenesis lead to an average increase of 29 de novo mutations per animal and 75% higher mutation rate per base pair in animals grown in crowded conditions compared to those grown in uncrowded conditions (Fig. 3). The numbers of de novo mutations and mutation rates seen in crowded animals could be underestimated due to the concurrent occurrence of purifying selection that removes deleterious variants within gene bodies during multiple generations of crowding selection (Fig. 4, Fig. 5, Fig. S3, and Fig. S4)^73^. Therefore, secreted CPR-4 induced by the crowding stress functions as a driver for generating genetic mutations, which facilitate adaptive changes and genome evolution to the crowding stress. Stress-induced mutagenesis has been well documented in bacteria^74–76^. However, whether and how stress-induced mutagenesis occurs in animals remain largely unclear^75,77^. Our findings fill the critical gaps in addressing this fundamental question and will have important implications in understanding adaptive evolution in diverse organisms.

After multiple generations of continuous growth in the crowded condition, animals exhibited biased mutation distributions in their genomes, with higher percentages of mutations detected in the intergenic regions and lower percentages of mutations found within gene bodies, compared with animals grown continuously in the uncrowded condition (Fig. 4 and Fig. S3). Loss of *cpr-4* eliminated biased mutation distributions in the genomes induced by persistent crowding stress, indicating that CPR-4-induced mutagenesis underlies biased mutation distributions in the genomes under crowding selection (Fig. 4 and Fig. S3). In this study, we show that persistent crowding stress and CPR-4 increased generation of heritable mutations, three of which characterized so far are all mutations in exons, suggesting that the lower percentages of mutations detected within gene bodies in animals grown in the crowded condition are unlikely due to lower mutagenesis rates in gene bodies. Importantly, those animals with heritable mutations in coding regions are all less competitive in growth, due to compromised gene functions, and likely will drop in numbers or disappear from the population under persistent crowding stress (Fig. 5d). Such negative selection ultimately contributes to biased mutation distributions towards the intergenic regions over the gene bodies. Therefore, CPR-4-induced mutagenesis underlies biased mutation distributions in the genomes through crowding selection, and potentially, facilitates adaptive genome evolution to persistent crowding stress.

As expected, animals that evolved from many generations of crowding stress showed a faster growth rate in the crowded condition than animals grown in the uncrowded environment (Fig. 6 and Fig. S5). Interestingly, both types of animals showed comparable growth rates in the uncrowded condition, indicating that the growth advantage observed in crowd-adaptive animals is only activated or induced by the crowded environment and is not permanent. Importantly, this inducible growth competitiveness of crowd-adaptive animals is also dependent on CPR-4, as loss of *cpr-4* prevented crowd-conditioned animals from gaining inducible growth advantage (Fig. 6 and Fig. S5). Again, because CPR-4 promotes generation of genome mutations and increase of mutation rates, which drive genome evolution upon many generations of crowding selection, these crowding-evolved genome changes, including biased mutation distributions, may enable animals to rapidly adapt to the crowded environment compared with animals evolved from the uncrowded condition and *cpr-4* mutant animals evolved from the same crowding selection, both of which lack CPR-4-mediated mutagenesis. This CPR-4-dependent, inducible growth advantage of crowd-adaptive animals is particularly interesting, because it enables animals to rapidly respond to the crowding stress, while maintaining the adaptive plasticity to cope with the potentially fluctuating environments in the early stage of evolution. This may represent an evolutionarily protective mechanism against environmental fluctuations before permanent adaptative changes become necessary^78,79^. Therefore, our study reveals a CPR-4-dependent mechanism crucial for adaptive plasticity, an evolutionary process whose underlying mechanisms remain poorly understood^78,80,81^.

The cathepsin B protease (CTSB) is highly conserved in the animal kingdom, with at least 45% sequence identity among cathepsin B proteins from different species (Fig. S6), which suggests conserved protein functions among the animals. As a lysosomal cysteine protease, CTSB has been shown to be secreted into the extracellular space under various stress conditions, such as radiation in *C. elegans*^31^ and chemotherapy and exercise in mammals^82–84^, or under pathological stress conditions, such as cancer, inflammation, and viral infection^83,85,86^, to exert important extracellular functions, including regulation of apoptosis, cell proliferation, cell motility and invasion, and neurogenesis^87^. Our findings uncover a new role of CTSB as a crucial responding factor and a driver of adaptive genome changes and evolution to the crowding stress in *C. elegans*, and potentially, in other organisms. Therefore, CTSB defines a new class of secreted signaling molecules, stresskines, that play crucial roles in mediating organisms’ responses to various stresses.

With an increasing world population projected to be at 8.1 billion in 2024, which is more than triple the number in 1950 (estimated at 2.5 billion), overpopulation and crowding stress have emerged as major challenges in contemporary societies, especially in the urban cities, where two-thirds of the world population live. Notably, the fertility rate of humans has decreased substantially, falling from 5 births per woman in 1950 to 2.3 births in 2021^88^. The negative impacts of overpopulation also include higher mortality rates and a high incidence of various neurological problems, such as depression and anxiety^89,90^. Therefore, our study may provide important molecular and mechanistic insights into the processes and underlying problems of high population density and facilitate future studies and treatments of vital human health issues associated with the escalating crowding stress.

## Online content

Methods, additional references, source data, extended data, supplementary information, acknowledgements, details of author contributions and competing interests, and statements of data and code availability are available at the online content.

## Methods

### Strains and culture conditions

*C. elegans* strains were cultured on 6-cm diameter nematode growth medium (NGM) plates with the *Escherichia coli* OP50 as the food source at room temperature using standard procedures^64^. We placed 600 µL of bacterial culture (∼ 0.7 OD600) on each NGM plate for uncrowded conditions. In crowded conditions, we used twice the amount of bacteria to accommodate the higher population density. The N2 Bristol strain was used as the wild-type strain. The following strains were used in this study: LGI, *hus-1(sm556*[*hus-1::Neongreen*])^32^, *daf-16(mu86)*; LGII, *smIs466(*P*cpr-4::cpr-4::flag)*^31^, *smIs564 (*P*skn-1::skn-1::gfp)*, *daf-22(m130)*; LGIII, *daf-2(e1370)*, *skn-1(zj15)*, *dvIs19 [P_gst-4_GFP]*; LGIV, *iaIs7 [P_nhr-57_GFP + unc-119(+)]*; LGV, *cpr-4(tm3718)*, *cpr-4(sm931)*, *akt-1(mg144)*; LGX, *pdk-1(sa680),* and single copy insertion of P*myo-2::mCherry*. Each mutant, transgene, and knock-in strain were backcrossed at least four times with N2 animals before being used in this study. Mice were maintained in a controlled environment with a 12-hour light/dark cycle (8:00 AM to 8:00 PM as the light period) and a constant room temperature of 23 ± 1°C. Both feed and water were provided ad libitum. Animal care and experiments were conducted in accordance with the guidelines for animal experiments at the University of Colorado. All experimental protocols were approved by the University of Colorado Institutional Animal Care and Use Committee and complied with the guidelines set forth by the US National Institutes of Health for the care and use of laboratory animals.

### Quantification of HUS-1::NeonGreen foci in germ cells

The gonad of *hus-1(sm556*[*hus-1::Neongreen*]) adult animals grown in the crowded or uncrowded condition was imaged with Z-stacks from the top to the bottom of the gonad using a Zeiss Axioplan 2 microscope equipped with a Cohu CCD camera and SlideBook 5 software (Intelligent Imaging Innovations, Inc., USA). The number of HUS-1::NeonGreen foci in the germline region from the mitotic zone to the mid-pachytene region was then quantified. For the uncrowded condition, each plate contained approximately 750 animals (Fig. 1c, d) or 1250 animals (Fig. 2). For the crowded condition, each plate contained approximately 3000 animals (Fig. 1c, d) or 5000 animals (Fig. 2).

### Generation of the *cpr-4* knockout mutant

The *cpr-4(sm931)* knockout mutant was generated using the CRISPR/Cas9 gene editing method^68^. An injection mix containing 50 ng/μL Cas9 expression construct pDD162, a *ben-1* sgRNA plasmid (45 ng/μL), two different *cpr-4* sgRNA plasmids (45 ng/μL each) targeting regions very close to the translation start and the stop codon of the *cpr-4* gene, respectively, and 20 ng/μL of oligonucleotide repair template was injected into young N2 adults. Injected animals were transferred to NGM plates with 14 μM benomyl (Sigma) and maintained at 25°C. Benomyl-resistant first generation (F1) animals were cloned out and screened for candidates that contained the desired deletion by polymerase chain reaction (PCR). F2 animals homozygous for the deletion were then isolated and confirmed by DNA sequencing. The oligonucleotide repair template: 5’ tgaagtaacttcagcatttgctctatcttgctatttgctcttttacaaaaataatctatttgattgaagtatttgttttcatacgtgtatatagatatga 3’.

### Collection of conditioned medium

Well-fed gravid adult animals were bleached and washed three times with the M9 buffer (40 mM Na_2_HPO_4_, 22 mM KH_2_PO_4_, 8.5 mM NaCl, and 18.7 mM NH_4_Cl). Their eggs were allowed to hatch and grow overnight in the M9 buffer without foods to obtain synchronized larval stage one (L1) animals. The number of synchronized L1 animals was quantified and the desired number of animals were placed on each NGM plate. For the crowded condition, approximately 1500, 2400, and 5400 animals, respectively, were placed in one NGM plate with a full bacterial lawn. As uncrowded controls for the crowded condition, the same numbers of animals were equally divided into four groups and placed in four NGM plates. At the larval stage 4 (L4), animals and bacteria were washed off the NGM plates with 2 ml M9 buffer and then spun down at 5500 rpm for 2 min. For the four uncrowded plates, the same 2 ml of M9 buffer were used to wash animals and bacteria off each of the four plates sequentially. The precipitated animals were placed back to a new NGM plate with a full bacterial lawn and continued their growth for another 16 hours. The supernatant was transferred to an eppendorf tube, adjusted to 2 ml final volume, and centrifuged at 12000 rpm for 10 min to remove the bacteria. The resulting supernatant was used as the conditioned medium, which was measured for its protein concentration and stored at −80℃ with 20% glycerol. The conditioned medium from animals grown for 16 hours after the L4 stage was collected similarly.

### Detection of secreted CPR-4::Flag by immunoblotting

Conditioned medium derived from P*cpr-4::cpr-4::flag; cpr-4(tm3718)* animals grown in uncrowded or crowded conditions was concentrated using a 10-kDa molecular mass cut-off centrifugal filter column (PALL) and its protein concentration was quantified. The concentrated conditioned medium (6 μg of total proteins in Fig. 1b or 1% volume in Fig. S1a, b) were resolved by a 12% SDS–PAGE gel and transferred to a PVDF membrane for immunoblotting. Secreted CPR-4::Flag was detected using a monoclonal antibody to the Flag epitope tag (1:2,000 dilution, Sigma) and a goat anti-mouse secondary antibody conjugated with horseradish peroxidase (1:10,000 dilution, Sigma).

### Assays for embryonic lethality, larval arrest, and brood size

In Fig. 1e-g and Fig. S1j-l, L4 stage *hus-1::neongreen* animals under uncrowded or crowded conditions with the indicated numbers of NeonGreen foci in their gonads, which were determined using a fluorescent Nomarski microscope, were rescued from the slides and placed in a NGM plate. After 2-hour recovery, animals were transferred to a new NGM plate for several different assays. For the embryonic lethality assays, 20-25 rescued gravid adults were allowed to lay eggs for 6 hours and then removed from the plates. After 24 hours, the number of eggs that did not hatch (scored as dead eggs, N_dead_) and the number of eggs that developed into larvae (N_live_) were scored and used to determine the rate of embryonic lethality [N_dead_/(N_dead_ + N_live_)].

For the larval arrest assays, 20 to 25 rescued gravid adults were allowed to lay eggs for 6 hours. The number of the larvae that hatched out from eggs was scored as the total number (T_L_). After 3 days, the number of animals that did not enter the L4 stage was scored (N_L1-L3_) and used to determine the rate of the larval arrest (N_L1-L3_/T_L_).

For the brood size assays, 10 to 15 rescued L4 larvae were individually placed on a single NGM plate. Every other day, the animals were transferred to new NGM plates until they died. The number of the eggs on each plate was scored and the total number of eggs laid by each individual animal was determined as the brood size of that animal.

For Fig. 2b-d and Fig. S2a-c, all three assays were conducted similarly, except that gravid adults or larvae were used directly for the assays, without the extra step to rescue them from the slides.

### Determination of SKN-1::GFP nuclear localization

The anterior intestines of young adult *smIs564 (Pskn-1::skn-1::gfp)* animals or *cpr-4(tm3718)*; *smIs564* animals grown under uncrowded (approximately 750 animals/plate) or crowded conditions (approximately 4,000 animals/plate) were imaged. An animal was scored as positive for SKN-1::GFP nuclear localization if all six nuclei in its first three pairs of intestinal cells exhibited nuclear SKN-1::GFP. The percentage of animals with SKN-1::GFP nuclear localization was determined by dividing the number of animals showing positive SKN-1 nuclear localization by the total number of animals scored.

### Quantification of *gst-4* and *nhr-57* expression levels using a GFP reporter

For the *gst-4* expression analysis, whole body fluorescent images of young adult *dvIs19 [Pgst-4GFP]* animals and *dvIs19; cpr-4(tm3718)* animals grown under uncrowded (approximately 750 animals/plate) or crowded conditions (approximately 4,000 animals/plate) were captured using a Zeiss LSM 800 laser scanning confocal microscope (Carl Zeiss). For *the nhr-*57 expression analysis, whole body fluorescent images of *iaIs7 [P_nhr-57_GFP + unc-119(+)]* animals grown in uncrowded (approximately 750 animals/plate) and crowded conditions (approximately 4,000 animals/plate), starting from the L1 larval stage until 16 hours after the L4 stage, or similar stage *iaIs7* animals in a hypoxic condition (0.5% oxygen for 5 hours) and in a normal, nonhypoxic condition were similarly captured, respectively. The whole body GFP fluorescence intensity in each animal was quantified using the Image J software with the measure parameter under the Analyze plugin. The mean GFP fluorescence intensity was used to determine the gene expression levels.

### Growing animals in the crowded or uncrowded condition for multiple generations

Single N2 animal, *cpr-4(tm3718)* animal, and N2 animal carrying the single-copy insertion of P*myo-2::mCherry* (cN2 as cherry-labeled N2) were placed in a NGM plate, respectively (See Fig. 3a and Fig. 6a). Their progeny (F1) were divided into two groups, one group for breeding in the uncrowded condition and the other in the crowded condition. For growth in the uncrowded condition, close to 20% of the F2 progeny (approximately 1000 animals) were transferred to a new NGM plate with a full bacterial lawn. Subsequently, the same number of animals were transferred to a new NGM plate each day for continuous propagation. For growth in the crowded condition, two-thirds of the F2 progeny (approximately 5000 animals) were transferred to a new NGM plate with a full bacterial lawn. Subsequently, the same number of animals on the plate were transferred to a new NGM plate twice a day for continuous propagation. In both conditions, animals always had sufficient OP50 food supplies. Three days of continuous breeding was considered as one generation and the breeding continued for multiple generations as indicated.

### Quantification of abnormal phenotypes

After 15 or 30 generations of continuous breeding in the crowded or uncrowded condition, 200 N2 or *cpr-4(tm3718)* animals (at the L4 or young adult stage) were randomly scored for abnormal phenotypes under a dissection microscope in each independent experiment. Animals with abnormal phenotypes were cloned out to individual NGM plates for further analysis to determine if the phenotype is heritable. The percentage of animals with abnormalities was determined from five independent experiments.

### Transgenic animals and rescue experiments

Transgenic animals were generated as described previously^91^. *exc-5-*containing fosmid WRM0622cA06 (at 20 ngμl^−1^) or *apa-2-*containing fosmid WRM0637dC06 (at 20 ngμl^−1^) was injected into *exc-5(sm775)* or *apa-2(sm780)* animals along with the pTG96 plasmid (at 20 ngμl^−1^), a co-injection marker containing a *sur-5::gfp* translational fusion, which is expressed in many cells and in most developmental stages^92^. As a negative control for rescue, pTG96 (at 20 ngμl^−1^) was injected alone into *exc-5(sm775)* or *apa-2(sm780)* animals. At least three independent transgenic lines were obtained and scored for each injection.

### Five-generation growth competition assays

In Fig. 6a-e and Fig. S5a-d, 50 L4 stage conditioned N2 or *cpr-4(tm3718)* animals were placed in the same NGM plate with 50 mCherry-labeled, conditioned N2 animals (cN2)(G0). The mixed animals were grown continuously in the uncrowded or crowded conditions for five more generations. For growth in the uncrowded condition, 20% of animals (approximately 1000 animals) on the G0 plate were transferred to a new NGM plate with a full bacterial lawn on day 3 and subsequently the same number of animals were transferred to a new NGM plate twice a day. For growth in the crowded condition, two-thirds of animals (approximately 5000 animals) on the G0 plate were transferred to a new NGM plate with a full bacterial lawn on day 3 and subsequently the same number of animals were transferred to a new NGM plate twice a day. In both conditions, animals always had sufficient OP50 food supplies. Three days of continuous breeding was considered as one generation and the breeding continued for five generations. At the first, third and fifth generation, one-third of the animals on the plate were mounted on a slide with 2% agarose pad and 300 L4 or young adult animals were randomly scored under the fluorescent dissection microscope. The number of cN2 animals among 300 animals was used to represent the percentage of cN2 animals in the population. In Fig. 5d, five-generation growth competition assays were conducted similarly, except that abnormal mutant animals and cN2 animals were used instead in the growth competition assays.

### Whole-genome sequencing analysis

The methods of whole-genome sequencing were used as described previously^93^. Multiple single L4 stage N2 or *cpr-4* mutant animals from 10 generations of continuous growth in the uncrowded and crowded conditions, respectively, were randomly isolated and propagated clonally for two or three generations. The animals were then collected and placed in standard worm lysis buffer [100 mM Tris-HCl (pH8.3), 500 mM KCl, 15 mM MgCl2, 0.1% Gelatin, and 1% SDS]. The genomic DNA was extracted by QIAGEN DNeasy kit, and the concentration and quality of purified DNA were determined by a Nanodrop 2000 spectrophotometer (Thermo Fisher Scientific) and electrophoresis. The genomic DNA was then fragmented into approximately 140 bp segments before library preparation. The genomic libraries were constructed by using automated Library Builder system (Thermo Fisher Scientific), and the sequencing templates were synthesized from the prepared libraries by the Ion Chef system according to the manufacturer’s protocol. The sequencing was completed by Ion Proton (Thermo Fisher Scientific) with the standard protocols (https://ioncommunity.thermofisher.com/docs/DOC-8775). Raw sequencing data were aligned to the *C. elegans* genome (version WS283) from WormBase (ftp://ftp.wormbase.org/pub/wormbase/), using the TMAP alignment program with the Burrows-Wheeler alignment algorithm (https://github.com/iontorrent/TS/tree/master/Analysis/TMAP). Variants were called using the Torrent Variant Caller (https://github.com/iontorrent/TS/tree/master/plugin/variantCaller) with default parameters to detect germ-line variants (https://github.com/iontorrent/TS/blob/master/plugin/variantCaller/pluginMedia/parameter_sets/amplis eq_germline_lowstringency_p1_parameters.json). For sequencing, the replicate numbers for each condition were indicated in figure legends. All samples had an average read depth of 20 or more. During the mapping process, an average of 0.57% of total reads per sample were discarded, primarily due to alignment with the *E. coli* genome. The total number of reads, discarded reads, bases with quality values greater than 20, and read depths relative to the *C. elegans* genome are summarized in Supplementary Table S1.

### Variants filtering and determination of de novo mutations and mutation rates

After initial variant calling, we applied a stringent custom filtering pipeline specifically to remove mutations that arise within sequences containing three or more consecutive identical nucleotides (e.g., AAA, TTT, CCC or GGG), as artifacts in homopolymeric regions are commonly seen in both Illumina and Ion Torrent sequencing^94,95^. After that, variants were further processed with post-calling filters using an in-house pipeline. The filtering methods are similar to those described previously^61,63,96^. Specifically, de novo mutations were called only if they met the following criteria:

1, Minimum of 5-fold coverage;

2, Base quality scores (Q scores) ≥25 for the candidate variants;

3, Coverage was not greater than 30-fold;

4, At least 20% of reads supporting the variant;

5, Variants were not reported in the corresponding F1 control;

6, Variants were not reported by any of the individual samples from the contrary group;

7, Substitutions excluded if located within or adjacent to an indel.

The mutation rate per base pair per generation was calculated by dividing the number of de novo mutations to the length of genome (100272607 bp; ver.WS283, downloaded from ftp://ftp.wormbase.org/pub/wormbase/) and the number of generations. De novo mutations were also categorized into four major types: deletion variants (DEL), single nucleotide polymorphism variants (SNV), multiple nucleotide polymorphism variants (MNV), and insertion variants (INS). SNVs were then subdivided into six categories based on specific nucleotide changes: T/A (A/T in reverse), T/C (A/G), T/G (A/C), C/A (G/T), C/T (G/A), and C/G (G/C). The number of mutations for each of these subgroups was determined similarly.

### Quantification of de novo mutations in different genomic regions

We annotated the genomic regions of de novo mutations using the reference of WBcel235 genome annotation (RefSeq.gtf from NCBI) with default output from Torrent Variant Caller. Mutations located in protein-coding regions or noncoding RNAs were classified as exonic, while those within introns or splice sites were classified as intronic. Both exonic and intronic mutations were collectively annotated as gene body mutations. Mutations not belonging to either of these categories were designated as intergenic. The numbers and percentages of mutations in intergenic regions and gene bodies, as well as in exons and introns were calculated.

### Collection of mouse plasma

A group of 20 6-week old C57BL/6 mice were initially housed together in cages for two weeks to facilitate their acclimation. Blood was collected from each mouse as Day 0 sample. The mice were then divided into two groups. One group were in cages with two mice per cage and the other group in five mice per cage. On Day 10 and Day 27, blood was collected from each mouse as Day 10 sample and Day 27 samples, respectively. To prepare plasma from blood, whole blood was mixed with heparin and centrifuged at 1,700 xg for 10 minutes. The plasma supernatant was collected and immediately placed on dry ice and stored at −80°C until analysis.

### Detection of secreted CTSB in mouse plasma

The mouse plasma were diluted 20 folds and the total protein level of each plasma was determined using the standard Bradford method (Bio-Rad). Equal amounts of total proteins (100 ng each) from the diluted mouse plasma were resolved by electrophoresis on 15% SDS–PAGE gels and transferred onto a PVDF membrane for immunoblotting analysis. CTSB was detected using a mouse monoclonal antibody to CTSB (1:2,000 dilution; ThermoFisher Catalog # 41-4800) and goat anti-mouse secondary antibodies conjugated with horseradish peroxidase (1:10,000 dilution; Sigma). The secreted CTSB levels were determined by quantifying the CTSB band intensities using the ImageJ software with the Analyzer plugin.

### Real-time quantitative RT-PCR analysis

Wild-type young adult animals grown under crowded (4000 animals/plate) and uncrowded (750 animals/plate) conditions, and as controls, grown in normal and hypoxic (0.5% oxygen for 5 hours) conditions, were collected, respectively, and subsequently lysed in TRI Reagent (Sigma, Cat # T9424) for total RNAs extraction following the standard protocol. Extracted total RNAs were used as templates for reverse transcription (RT) to synthesize the first strand cDNA using the SuperScript III First Strand Synthesis System (Invitrogen, Cat # 18080051). Quantitative PCR analysis was conducted using Applied Biosystem 7500 Fast Real-Time PCR System and the Power SYBR Green PCR Master Mix (Applied Biosystem, Cat # 4367659). Each PCR reaction contained 400 ng cDNA in a final volume of 20 μL. Amplifications were performed in MicroAmp Fast 96-Well Reaction Plate (Applied Biosystems, Cat # 4346907) placed in the 96-well, real-time PCR detection system. Melting curve analysis was performed after the final PCR cycle to examine the specificity of primers in each reaction. PCR reactions were performed in quadruplicate for each condition and target gene. The transcription of the *rpl-26* gene was used as the reference. The data were analyzed by the Livak method. The primers for *csyl-1* are 5’ GAAACACTGGAATCGCATTGG 3’ (forward) and 5’ TGTCCCTTTGGTTTGTCTCC 3’ (reverse). The primers for *egl-9* are 5’ TCCACAGCAACTATTCCACC 3’ (forward) and 5’ GGAGTTAGAAGGAGTTGGTCG 3’ (reverse). The primers for *hif-1* are 5’ TGGACAGGGTACAACTAATGC 3’ (forward) and 5’ GTTTGAATCCAGGCAAGTGTG 3’ (reverse). The primers for *rhy-1* are 5’ CCGTTCCAGTCTCATTGTCG 3’ (forward) and 5’ GATGAATTGGTGAAGTGGCAG 3’ (reverse). The primers for *rpl-26* are 5’ ATGAAGGTCAATCCGTTCGT 3’ (forward) and 5’ AGGACACGTCCAGTGTTTCC 3’ (reverse).

### Confocal microscopy and image analysis

Images in Fig. 1c and Fig. 2e were taken with a 63× objective and images in Fig. S1e and f and Fig. S2e were captured with a 10× objective, using a Zeiss LSM 800 laser scanning confocal microscope (Carl Zeiss). Images in Fig. 5b and Fig. S4d,e were captured with a 20x objective using a Zeiss Axioplan 2 microscope (Carl Zeiss) equipped with a Cohu CCD camera and SlideBook 5 software (Intelligent Imaging Innovations, Inc., USA).

### Statistical analysis

The statistical analysis was performed using Student’s *t*-test, two-sample *z*-test, Wilcoxon rank-sum test, multinomial logistic regression followed by the Dunnett-type comparisons test or binomial generalized linear model (logit) followed by the Dunnett-type comparisons test, as indicated in the figure legends, using the software Prism 9 or R.

### Data Availability

The whole genome sequencing data generated in this study have been deposited in the NCBI database and are available under BioProject PRJNA1118417 (http://www.ncbi.nlm.nih.gov/bioproject/1118417). Codes used in this study can be accessed from https://github.com/Anti-Bin/crowding-mutation-analysis. Source data are provided with this paper. Experimental materials are available upon requests to D.X.

## Acknowledgements

We thank members of the Xue lab for discussions, V. Zaberezhnyy for help with the mouse work, and D. Stock for help with some of the experiments. This work is supported in part by NIH grant R35 GM118188 (to D.X.), the Veteran’s Administration grant 1 I01 BX004495 (to J.D.), NIGMS/NIH grant 5T32GM141742-02 (to B.J.), the National Basic Research Program of China grant 2019YFA0508401 (to G.O.), and the China Postdoctoral Science Foundation grant 2022M711844 (to D.L.).

## Author contributions

B.Y. and D.X. conceived the study, analyzed the data, and wrote the paper. B.Y. performed most of the experiments and bioinformatics analysis in this study. Y.S. and S.M. performed the whole genome sequencing analysis. B.J. and J.D. performed mouse crowding experiments. E.L., D.L., G.O., J.J., and Y.H. assisted in some of the experiments and data collection. All authors read and edited the manuscript.

---

**Fig. S1.**
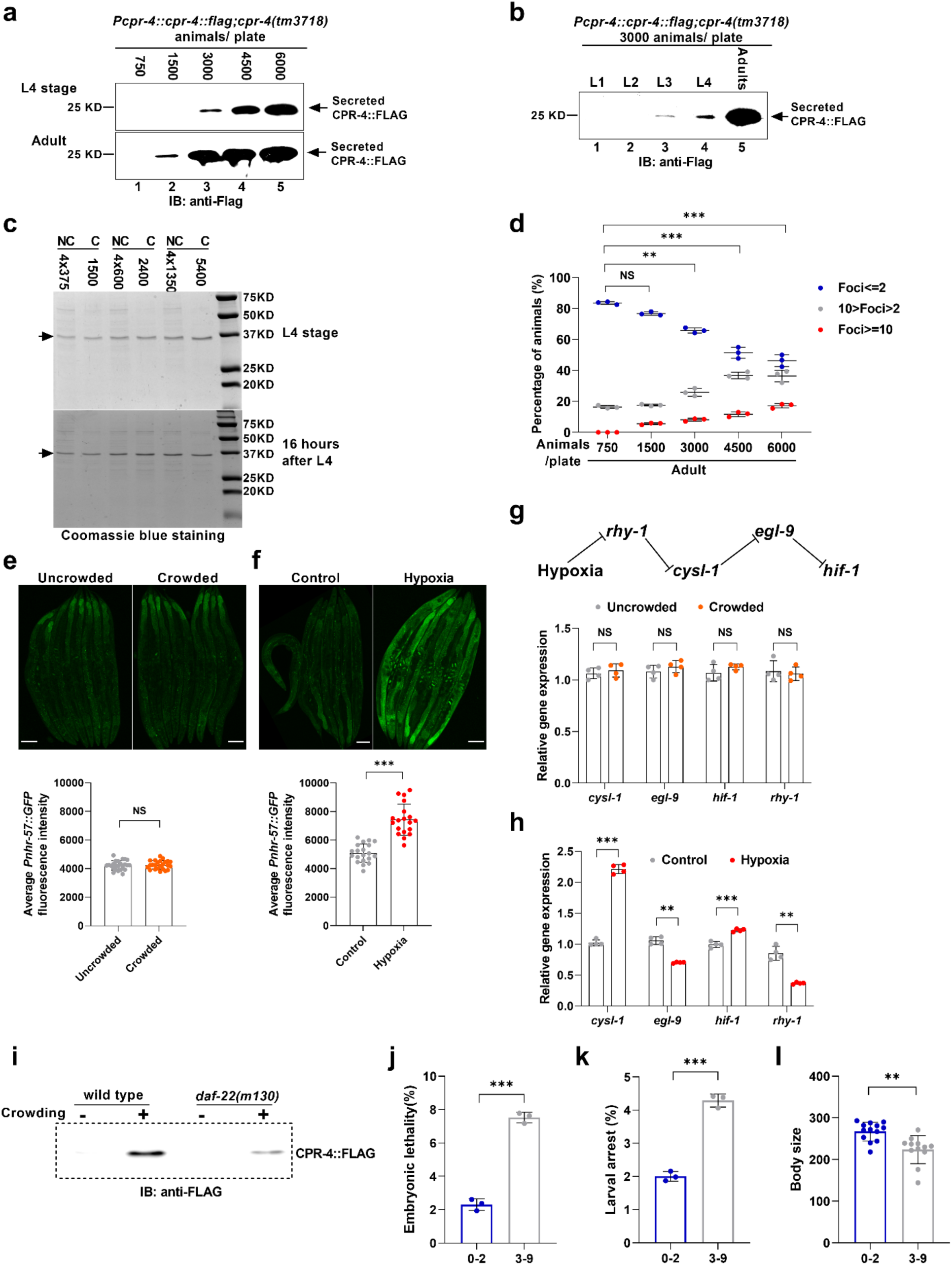
CPR-4 secretion is positively correlated with the population density and germ cell DNA damage in *C. elegans*. **a**, **b**, CPR-4::FLAG secretion from P*cpr-4::cpr-4::flag*; *cpr-4(tm3718*) animals grown at the indicated density was examined at the L4 stage and 16 hours after the L4 stage (**a**) and in five different developmental stages (**b**). Conditioned medium was collected as described in Materials and Methods and adjusted to the same volume. One percent of the conditioned medium were resolved in 12% SDS polyacrylamide gel (SDS-PAGE) and detected by an antibody to the FLAG epitope using immunoblotting (IB). **c**, Equal amounts (6 μg total protein each) of noncrowded (NC) and crowded (C) conditioned medium as indicated were resolved by 12% SDS-PAGE, which was stained by Coomassie Blue. The strong protein band of unknown identity (indicated by an arrow) was used as the loading control. **d**, Percentages of *hus-1::neongreen* knockin animals with the indicated numbers of NeonGreen foci in their gonads are shown. Animals were grown at the indicated density and examined 16 hours after the L4 stage. Three biologically independent experiments were performed for each density. The total number of animals analyzed from left to right are 103, 109, 111, 119, and 106. **e**, **f**, Representative fluorescent images of P*nhr-57::gfp* transgenic animals grown in uncrowded and crowded conditions (**e**), starting from the L1 larval stage until 16 hours after the L4 stage, or similar stage P*nhr-57::gfp* transgenic animals in a normal, nonhypoxic condition and a hypoxic condition (0.5% oxygen for 5 hours)(**f**), respectively, are shown. The average fluorescence intensity of P*nhr-57::gfp* animals in these four different conditions was quantified and shown (bottom panels). n ≥20 animals examined for each condition in each of three biologically independent experiments. Scale bars, 100 μm. **g**, **h**, The expression levels of four hypoxia-related genes, *hif-1*, *rhy-1*, *egl-9*, and *cysl-1*, are altered significantly in animals under the hypoxia condition, compared with those under the nonhypoxic condition (**h**), but are comparable in uncrowded and crowded animals (**g**). n=4 biologically independent experiments were conducted for each condition. The genetic relationship among the hypoxia-related genes under the hypoxic condition is shown in **g**. **i**, CPR-4::FLAG secretion from *Pcpr-4::cpr-4::flag; cpr-4(tm3718)* animals and *Pcpr-4::cpr-4::flag daf-22(m130); cpr-4(tm3718)* animals grown in one NGM plate (crowded, 4500 animals/plate) or evenly distributed in four NGM plates (noncrowded, 1125 animals/plate) was examined at 16 hours after the L4 stage. Conditioned medium was collected as in **a** and adjusted to the same volume. One percent of the conditioned medium were resolved in 12% SDS-PAGE and detected by an antibody to the FLAG epitope using immunoblotting. **j**-**l**, The percentages of embryonic lethality (**j**) and larval arrest (**k**) observed in progeny and the brood sizes (**l**) of *hus-1::neongreen* adult animals grown in uncrowded conditions with the indicated number of NeonGreen foci, which were rescued from agarose pads after microscopy screens. Three biologically independent experiments in **j** and **k**. The total numbers of adult animals analyzed from left to right are 60 and 60 in **j** and **k**, 13 and 12 in **l**, with the foci numbers from one gonad arm. Total numbers of embryos scored in **j**: 3288 (foci≤2) and 2125 (foci between 3-9). Total numbers of larvae scored in **k**: 3211 (foci≤2) and 1990 (foci between 3-9). Data mean ± s.d.. NS, not significant, \*\**P* < 0.01, \*\*\**P* < 0.001, two-sided *z*-test in **d** and two-sided, unpaired *t*-test in **e**-**k**.

**Fig. S2.**
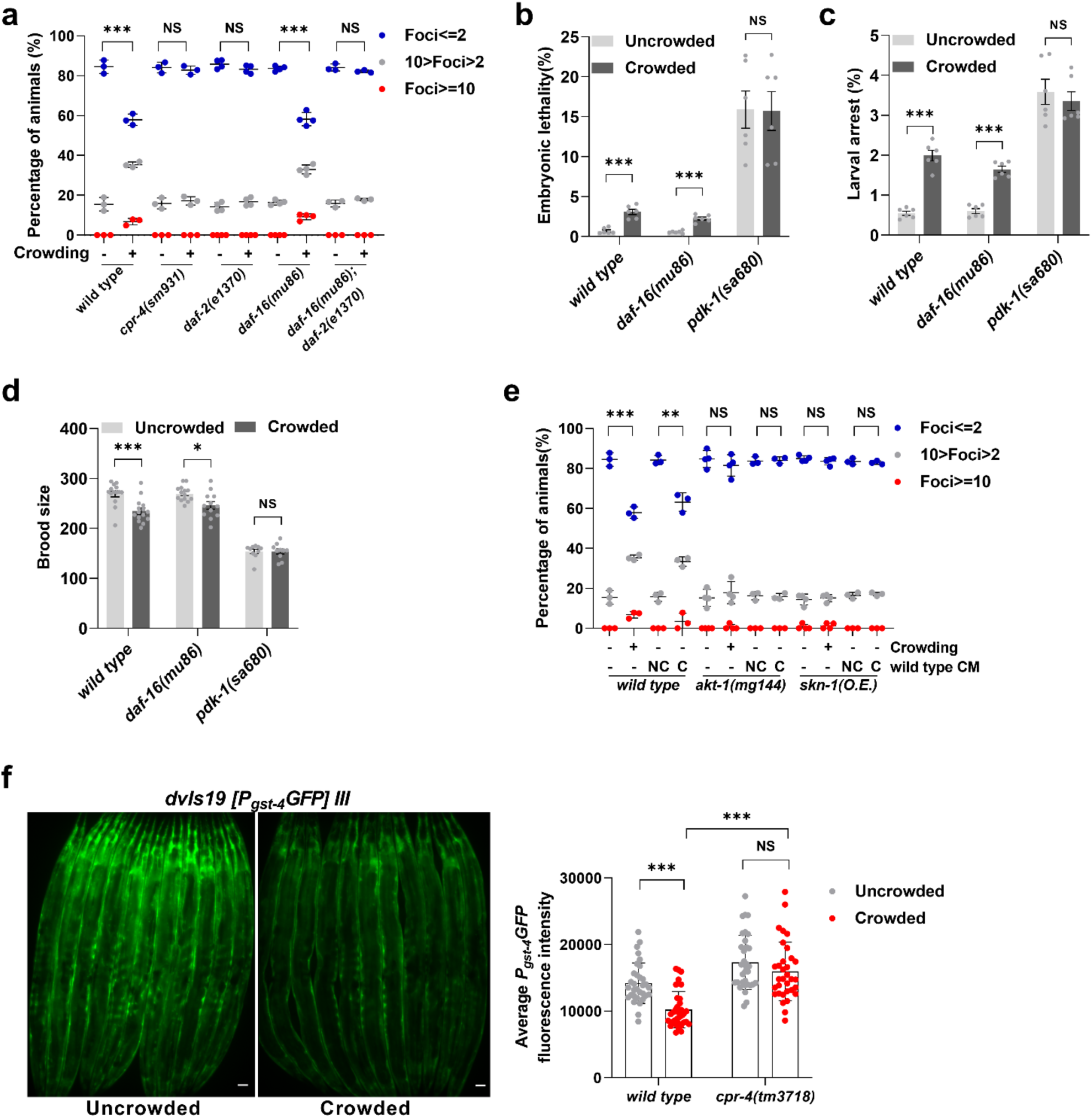
Crowding responses are DAF-16 independent. **a**,**e**, Percentages of the *hus-1::neongreen* adult animals with the indicated genotype and grown in uncrowded or crowded conditions (**a**), or in uncrowded conditions and then treated with NC-CM or C-CM (**e**), showing gonad NeonGreen foci ≤2 (blue), between 3-9 (grey), and ≥10 (red), respectively. n ≥33 gonads examined in each of at least three biologically independent experiments. **b-d**, Percentages of embryonic lethality (**b**) and larval arrest (**c**) observed in progeny and brood sizes (**d**) observed in adult animals with the indicated genotype, which were grown in the uncrowded and crowded conditions, respectively. Six biologically independent experiments were conducted in each condition and genotype. **f**, *gst-4* expression is downregulated in wild-type animals but not in *cpr-4(tm3718)* animals in the crowded condition, shown by fluorescent images of the *P_gst-4_gfp* transgenic animals grown in uncrowded and crowded conditions, respectively (left panel), and by their average fluorescence intensity in both conditions (right panel, *n* = 33). Scale bars, 50 μm. The total numbers of adult animals analyzed from left to right are 117, 119, 94, 99, 144, 174, 153, 164, 107, and 112 in **a**, 120, 120, 120, 120, 120, and 120 in **b** and **c**, 14, 15, 14, 12, 10, and 10 in **d,** 117, 119, 114, 114, 158, 157, 99, 101, 159, 158, 85, and 88 in **e**, and 32, 33, 34, and 33 in **f**. Total numbers of embryos scored in **b**: 4064, 3758, 4885, 4361, 3726, and 4139 from left to right. Total numbers of larvae scored in **c**: 4038, 3653, 4887, 4336, 3225, and 3592 from left to right. Data are mean ± s.d. (**a, d-f)** and mean ± s.e.m (**b**, **c**). NS, not significant, \**P* < 0.05, \*\**P* < 0.01, \*\*\**P* < 0.001, two-sided, unpaired *t*-test (**b-d, f)** and two-sided *z*-test (**a**, **e**).

**Fig. S3.**
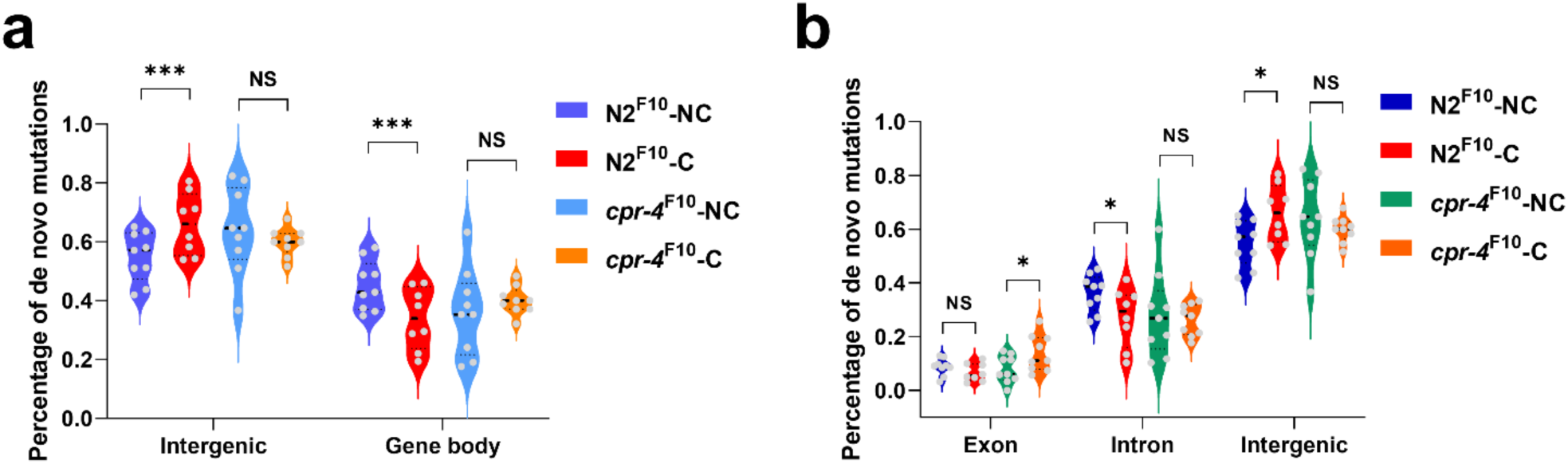
Persistent crowding stress and CPR-4 cause biased mutation distributions in genomes of crowded animals. **a**, **b**, Percentages of de novo mutations identified in the intergenic regions and gene bodies (**a**) or in exons, introns, and intergenic regions (**b**) in N2 and *cpr-4(tm3718)* animals at the 10^th^ generation of continuous growth in crowded (C) and uncrowded (NC) conditions, respectively. n=9 (N2^F10^-NC), 8 (N2^F10^-C), 9 (*cpr-4*^F10^-NC), and 9 (*cpr-4*^F10^-C) from two biologically independent experiments. NS, not significant, \**P* < 0.05, \*\*\**P* < 0.001, binomial generalized linear model (logit) followed by the Dunnett-type comparisons test (**a**) or multinomial logistic regression followed by the Dunnett-type comparisons test (**b**).

**Fig. S4.**
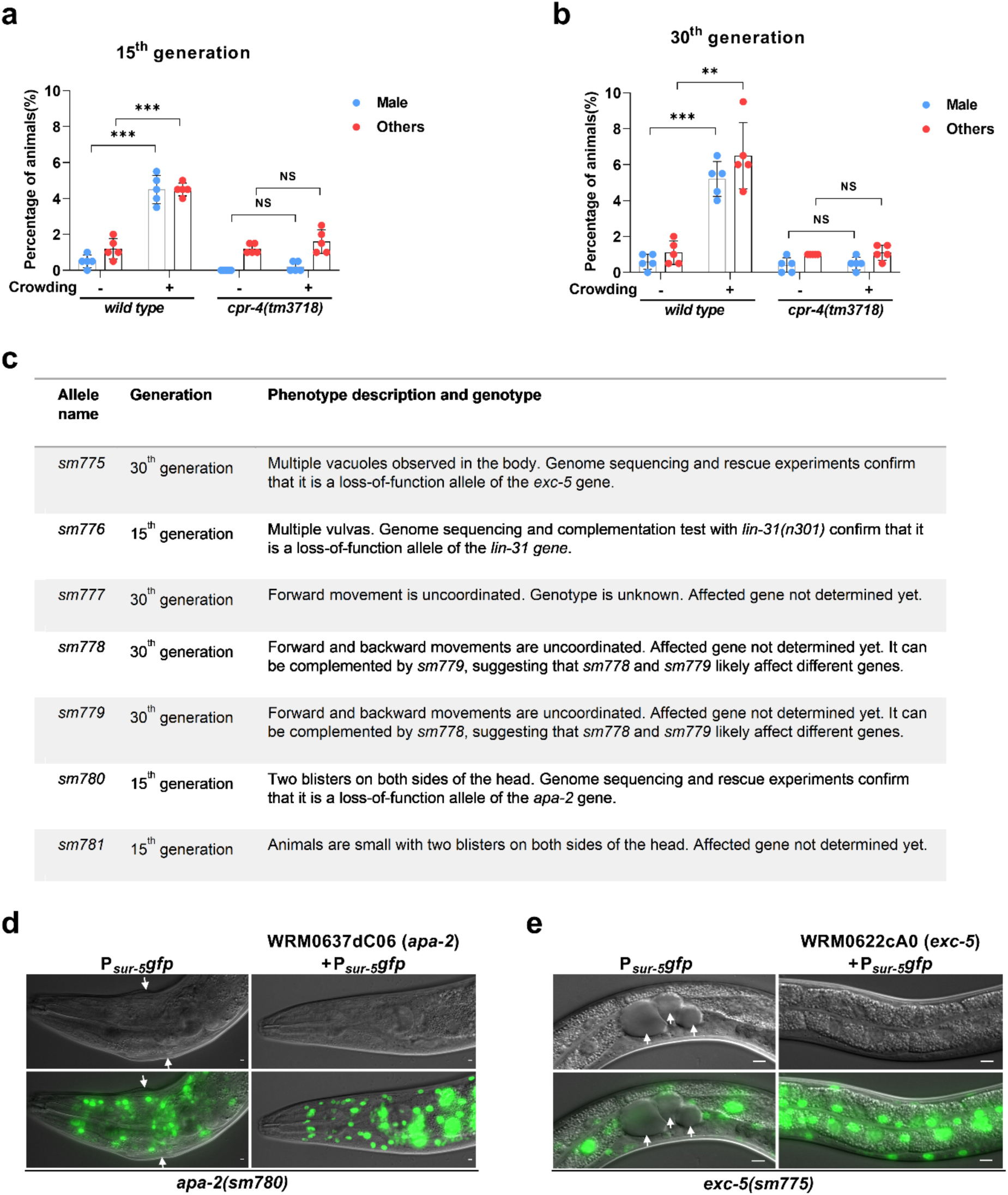
Persistent crowding stress and *cpr-4* increase generation of heritable mutations. **a**, **b**, Percentages of N2 (wild-type) and *cpr-4(tm3718)* animals that were males or exhibited other abnormal phenotypes (others) under continuously crowded and uncrowded conditions scored at the 15^th^ and the 30^th^ generation (each condition with five biologically independent experiments). Total numbers of animals scored are 1000 in each condition. **c**, Heritable mutants identified from the wild-type populations grown in continuously crowded conditions at the 15^th^ or the 30^th^ generation, their mutant phenotypes, and genes affected, if determined. **d**, **e**, Representative differential interference contrast (DIC) and fluorescent images of *apa-2(sm780)* transgenic animals (**d**) and *exc-5(sm775)* transgenic animals (**e**) are shown. The blister defect of *apa-2(sm780)* animals (**d**) and the vacuolar defect of *exc-5(sm775)* animals (**e**) are indicated with arrows and are rescued by the fosmid WRM0637dC06 containing the *apa-2* coding region and the fosmid WRM0622cA06 containing the *exc-5* coding region, respectively. P*_sur-5_gfp* is the transgenic marker used, which drives GFP expression in all cells in most developmental stages. Data are mean ± s.d. (**a**, **b**). NS, not significant, \*\**P* < 0.01, \*\*\**P* < 0.001, two-sided, unpaired *t*-test.

**Fig. S5.**
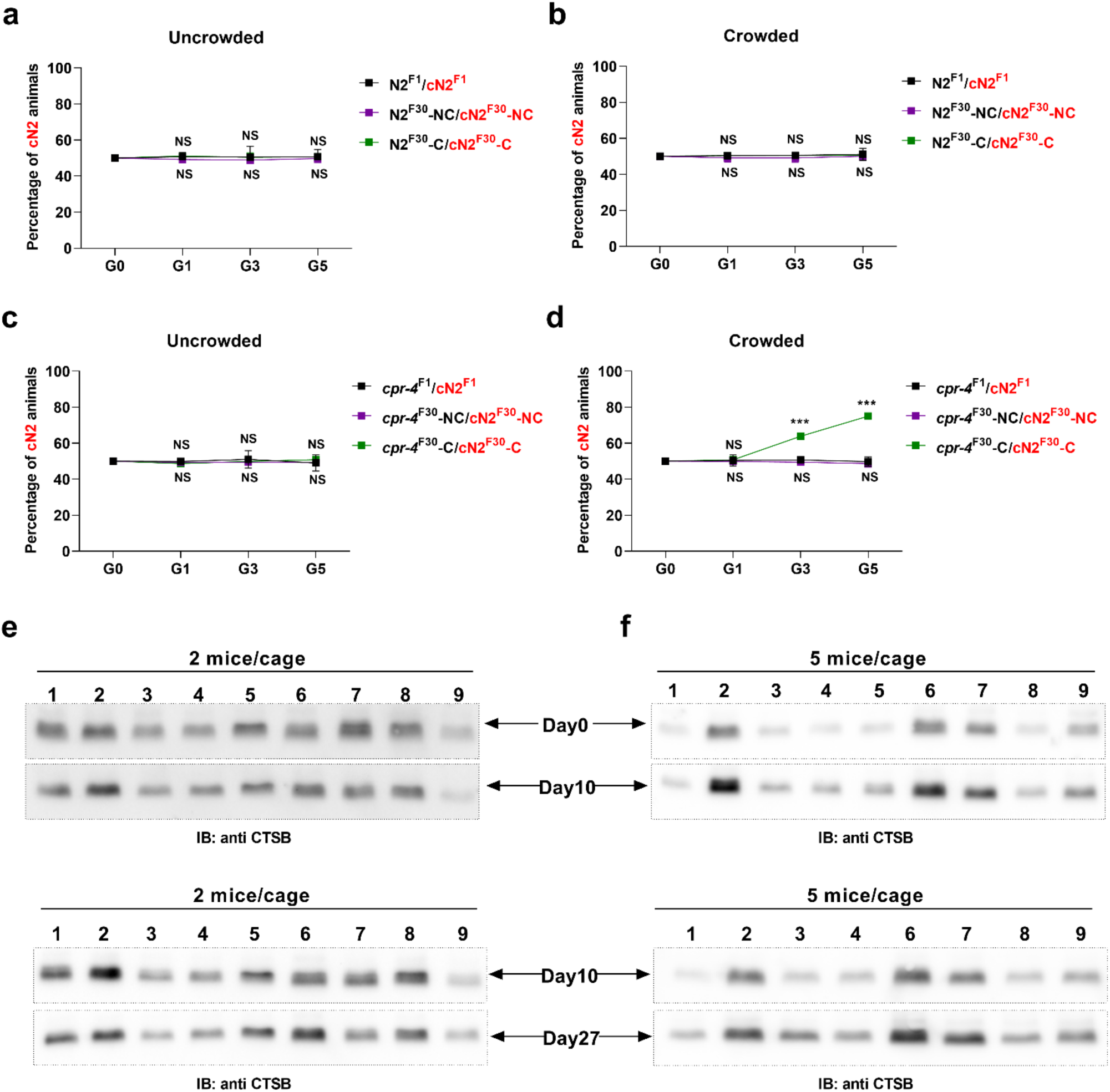
*cpr-4* promotes inducible growth advantage after 30 generations of continuous crowding stress. **a-d**, Five-generation growth competition assays between the indicated groups of animals grown in uncrowded (**a**, **c**) and crowded (**b**, **d**) conditions as illustrated in Fig. 6a. One group of N2 animals were labeled by mCherry (cN2 in red) expressed from the P*_myo-2_*mCherry single-copy integrated transgene. The percentages of mCherry-labeled animals in the populations at the first (G1), third (G3), and fifth (G5) generations were scored (*n* = 300 in each experiment). First generation (F1) animals shown in Fig. 6a were used as unconditioned controls. Animals at the 30th generation (F30) grown in either crowded (C) or uncrowded (NC) conditions were used in the growth competition assays. Three biologically independent experiments were performed for each pair of animals and condition. **e**, **f**, Immunoblotting images showing the levels of secreted forms of CTSB in plasma from mice housed in 2 mice per cage (**e**) and 5 mice per cage (**f**) on Day 0 (before crowd conditioning), Day 10, and Day 27. Plasma with equal amounts of total protein (100 ng) were resolved in 15% SDS polyacrylamide gels (SDS-PAGE) and detected by a monoclonal antibody to the human CTSB protein using immunoblotting (IB). 8-9 mice were examined for each group. Plasma from the same mouse at three different time points (indicated with the same mouse number) were analyzed. #5 mouse in 5 mice/cage was lost on Day 27. Data are mean ± s.d. (**a**, **b**, **c**, and **d**). NS, not significant, \*\*\**P* < 0.001, binomial generalized linear model (logit) followed by the Dunnett-type comparisons test.

**Fig. S6.**
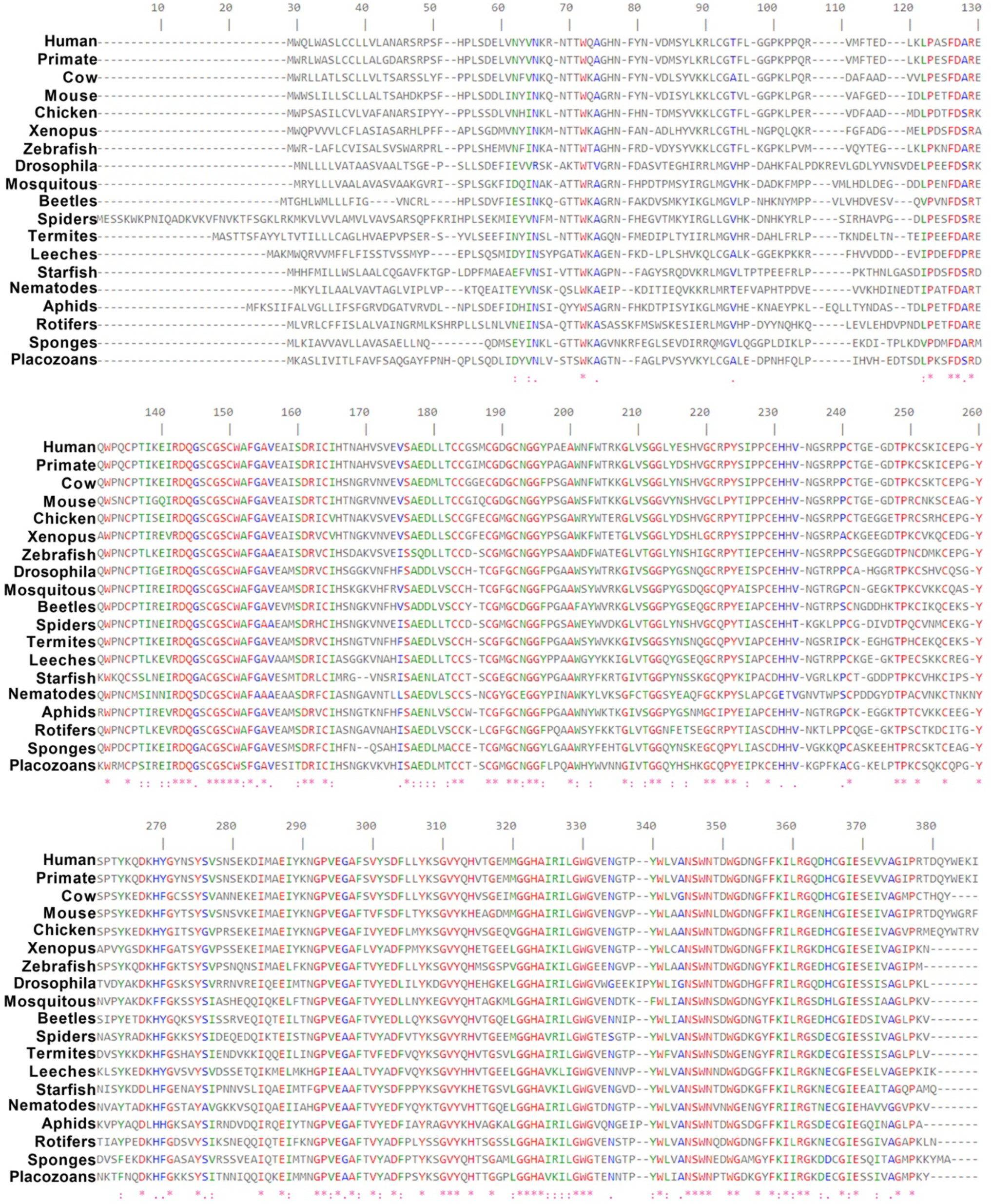
Cathepsin B protein is highly conserved in animals. Sequence alignment of Cathepsin B proteins from 19 different animal species, including human, primate (*Macaca mulatta*), cow, mouse, chicken, Xenopus, Zebrafish, *Drosophila melanogaster*, Mosquitos (*Culex quinquefasciatus*), beetles (*Sitophilus oryzae*), spider (*Araneus ventricosus*), termite (*Zootermopsis nevadensis*), leech (*Helobdella robusta*), starfish (*Acanthaster planci*), nematode (*C. elegans*), Aphid (*Acyrthosiphon pisum*), Rotifer (*Adineta steineri*), Sponge (*Suberites domuncula*), and Placozoan (*Trichoplax adhaerens*). Residues that are identical in all Cathepsin B proteins are highlighted in red, residues that are identical or highly similar are highlighted in green, and residues that are identical or weakly similar are highlighted in blue.

**Table S1.** Additional information on whole-genome sequencing analysis.

| **Sample** | **Total reads** | **Mapped reads** | **Discarded reads** | **>Q20 (bases)** | **Depth** |
| --- | --- | --- | --- | --- | --- |
| N2 10th uncrowded 1 | 19048804 | 18943326 | 105478 | 2,619,140,474 | 28.47 |
| N2 10th uncrowded 2 | 18144502 | 18030706 | 113796 | 2,474,813,033 | 26.88 |
| N2 10th uncrowded 3 | 16051967 | 15943883 | 108084 | 2,198,638,340 | 24.04 |
| N2 10th uncrowded 4 | 14645043 | 14574500 | 70543 | 2,036,685,613 | 22.13 |
| N2 10th uncrowded 5 | 16996912 | 16896584 | 100328 | 2,304,614,962 | 24.88 |
| N2 10th uncrowded 6 | 14056689 | 13979221 | 77468 | 1,929,305,890 | 20.53 |
| N2 10th uncrowded 2nd 1 | 15729265 | 14182513 | 1546752 | 2,194,335,609 | 21.53 |
| N2 10th uncrowded 2nd 2 | 15688670 | 14193025 | 1495645 | 2,184,249,754 | 21.31 |
| N2 10th uncrowded 2nd 3 | 17580293 | 15827261 | 1753032 | 2,393,470,501 | 23.11 |
| N2 10th crowded 1 | 18808836 | 18708672 | 100164 | 2,520,575,129 | 26.72 |
| N2 10th crowded 2 | 17543967 | 17462583 | 81384 | 2,373,712,811 | 25.44 |
| N2 10th crowded 3 | 19614248 | 19511057 | 103191 | 2,626,954,116 | 28.18 |
| N2 10th crowded 4 | 19422452 | 19323980 | 98472 | 2,592,401,671 | 27.84 |
| N2 10th crowded 5 | 17965472 | 17874448 | 91024 | 2,382,563,821 | 25.72 |
| N2 10th crowded 2nd 1 | 18278347 | 16121086 | 2157261 | 2,465,684,196 | 23.1 |
| N2 10th crowded 2nd 2 | 18342627 | 16225493 | 2117134 | 2,468,556,850 | 23.18 |
| N2 10th crowded 2nd 3 | 17996124 | 15995590 | 2000534 | 2,438,982,076 | 23.11 |
| *cpr-4* 10th uncrowded 1 | 16603478 | 16506066 | 97412 | 2,269,415,330 | 23.87 |
| *cpr-4* 10th uncrowded 2 | 17368372 | 17295300 | 73072 | 2,444,921,863 | 26.03 |
| *cpr-4* 10th uncrowded 3 | 16526986 | 16427339 | 99647 | 2,311,310,605 | 24.66 |
| *cpr-4* 10th uncrowded 2nd 1 | 22937705 | 20778747 | 2158958 | 3,137,079,606 | 31.76 |
| *cpr-4* 10th uncrowded 2nd 2 | 14127265 | 12571932 | 1555333 | 1,879,878,559 | 17.79 |
| *cpr-4* 10th uncrowded 2nd 3 | 14953888 | 13470856 | 1483032 | 2,051,009,769 | 19.84 |
| *cpr-4* 10th uncrowded 2nd 4 | 13243208 | 11920299 | 1322909 | 1,601,113,198 | 15.07 |
| *cpr-4* 10th uncrowded 2nd 5 | 13454005 | 11933307 | 1520698 | 1,780,765,085 | 16.82 |
| *cpr-4* 10th uncrowded 2nd 6 | 16071487 | 14424129 | 1647358 | 2,214,816,588 | 21.3 |
| *cpr-4* 10th crowded 1 | 14272943 | 14183637 | 89306 | 1,978,610,732 | 21.15 |
| *cpr-4* 10th crowded 2 | 19397685 | 19243401 | 154284 | 2,670,441,844 | 28.4 |
| *cpr-4* 10th crowded 3 | 18455693 | 18324143 | 131550 | 2,557,773,118 | 27.31 |
| *cpr-4* 10th crowded 4 | 15444430 | 15348672 | 95758 | 2,153,142,555 | 22.88 |
| *cpr-4* 10th crowded 5 | 15864969 | 15778359 | 86610 | 2,201,264,394 | 23.36 |
| *cpr-4* 10th crowded 2nd 1 | 12282593 | 10891912 | 1390681 | 1,608,666,750 | 15.08 |
| *cpr-4* 10th crowded 2nd 2 | 13117040 | 11842723 | 1274317 | 1,780,484,602 | 17.05 |
| *cpr-4* 10th crowded 2nd 3 | 14110007 | 12572775 | 1537232 | 1,853,330,278 | 17.39 |
| *cpr-4* 10th crowded 2nd 4 | 18952639 | 16952144 | 2000495 | 2,597,823,105 | 24.64 |
| *sm778* | 20274553 | 17875490 | 2399063 | 3,276,281,305 | 25.35 |
| *sm779* | 21353242 | 19571422 | 1781820 | 3,453,228,624 | 28.12 |
| *sm777* | 19565548 | 19033662 | 531886 | 3,129,288,678 | 27.44 |
| *sm775* | 22309644 | 22136777 | 172867 | 3,577,248,242 | 32.4 |
| *sm780* | 18971458 | 18664441 | 307017 | 3,121,632,479 | 27.97 |
| *sm781* | 19918752 | 14239287 | 5679465 | 3,328,226,275 | 19.76 |
| *sm776* | 17046827 | 16588920 | 457907 | 2,763,212,799 | 24.33 |
| N2_control | 22200595 | 22110159 | 90436 | 3,639,932,910 | 33.31 |

**Table S2.**
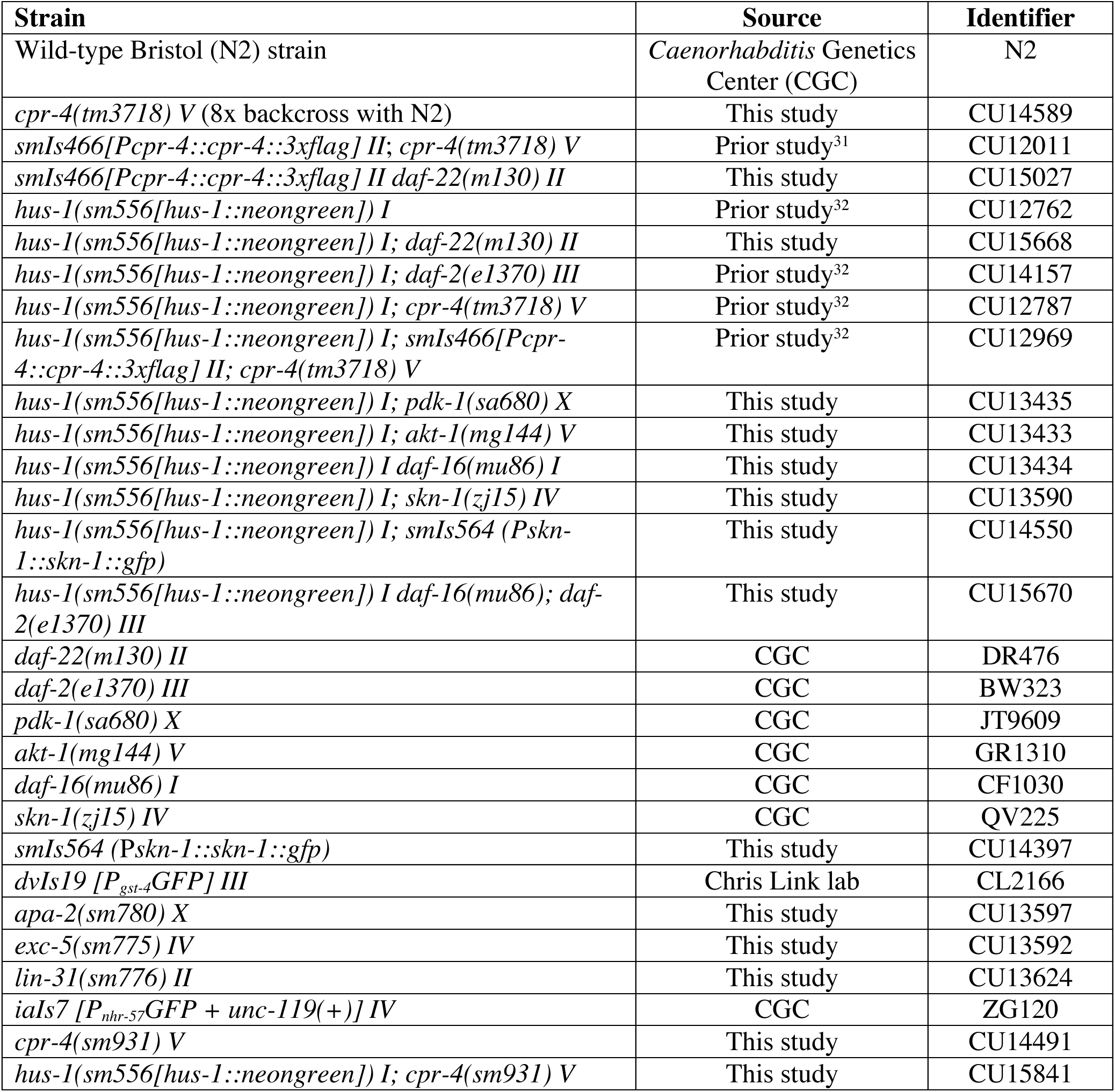
*C. elegans* strain information.

